# LAMP-Seq: Population-Scale COVID-19 Diagnostics Using Combinatorial Barcoding

**DOI:** 10.1101/2020.04.06.025635

**Authors:** Jonathan L. Schmid-Burgk, Ricarda M. Schmithausen, David Li, Ronja Hollstein, Amir Ben-Shmuel, Ofir Israeli, Shay Weiss, Nir Paran, Gero Wilbring, Jana Liebing, David Feldman, Mikołaj Słabicki, Bärbel Lippke, Esther Sib, Jacob Borrajo, Jonathan Strecker, Julia Reinhardt, Per Hoffmann, Brian Cleary, Michael Hölzel, Markus M. Nöthen, Martin Exner, Kerstin U. Ludwig, Aviv Regev, Feng Zhang

**Affiliations:** Broad Institute of MIT and Harvard Cambridge, MA 02142, USA; McGovern Institute for Brain Research, Massachusetts Institute of Technology, Cambridge, MA 02139, USA; Department of Brain and Cognitive Sciences, Massachusetts Institute of Technology, Cambridge, MA 02139, USA; Department of Biological Engineering, Massachusetts Institute of Technology, Cambridge, MA 02139, USA; Department of Biology, Massachusetts Institute of Technology, Cambridge, MA 02139, USA; Klarman Cell Observatory, Massachusetts Institute of Technology, Cambridge, MA 02139, USA; Koch Institute for Integrative Cancer Research, Massachusetts Institute of Technology, Cambridge, MA 02139, USA; Institute of Clinical Chemistry and Clinical Pharmacology, University Hospital Bonn, 53127 Bonn, Germany; Institute of Hygiene and Public Health, University Hospital Bonn, 53127 Bonn, Germany; Institute of Human Genetics, University Hospital Bonn, 53127 Bonn, Germany; Institute for Experimental Oncology, University Hospital Bonn, 53127 Bonn, Germany; Department of Infectious Diseases, Israel Institute for Biological Research, Ness Ziona, Israel; Department of Biochemistry and Molecular Genetics, Israel Institute for Biological Research, Ness Ziona, Israel; Department of Biochemistry and Institute for Protein Design, University of Washington, Seattle, WA 98195, USA; Department of Medical Oncology, Dana-Farber Cancer Institute, Boston, MA 02215, USA; Division of Translational Medical Oncology, German Cancer Research Center (DKFZ) and National Center for Tumor Diseases (NCT), 69120 Heidelberg, Germany; Genomics Research Group, Department of Biomedicine, University of Basel, Switzerland; Howard Hughes Medical Institute, Cambridge, MA 02139, USA

## Abstract

The ongoing SARS-CoV-2 pandemic has already caused devastating losses. Exponential spread can be slowed by social distancing and population-wide isolation measures, but those place a tremendous burden on society, and, once lifted, exponential spread can re-emerge. Regular population-scale testing, combined with contact tracing and case isolation, should help break the cycle of transmission, but current detection strategies are not capable of such large-scale processing. Here we present a protocol for LAMP-Seq, a barcoded Reverse-Transcription Loop-mediated Isothermal Amplification (RT-LAMP) method that is highly scalable. Individual samples are stabilized, inactivated, and amplified in three isothermal heat steps, generating barcoded amplicons that can be pooled and analyzed *en masse* by sequencing. Using unique barcode combinations per sample from a compressed barcode space enables extensive pooling, potentially further reducing cost and simplifying logistics. We validated LAMP-Seq on 28 clinical samples, empirically optimized the protocol and barcode design, and performed initial safety evaluation. Relying on world-wide infrastructure for next-generation sequencing, and in the context of population-wide sample collection, LAMP-Seq could be scaled to analyze millions of samples per day.

## Introduction

As of June 2020, the global spread of a novel coronavirus, SARS-CoV-2, has resulted in over 6,700,000 confirmed cases and 393,000 deaths (Johns Hopkins CSEE Covid tracker (Dong et al., 2020)). Early epidemiological studies indicate that the exponential spread of COVID-19, the disease caused by SARS-CoV-2, can be slowed by restrictive isolation measures (Chinazzi et al., 2020), but these measures place an enormous burden on societies and economies. Moreover, once isolation measures are lifted, exponential spread is predicted to resume (Li et al., 2020). At the same time, many infected individuals do not show any symptoms, remain untested, and thereby unknowingly contribute to the spread of infection (Yu et al., 2020). Repeated population-scale testing that enables identification of all infected individuals regardless of their symptomatic or contact status was predicted as an effective measure to help combat the transmission of SARS-CoV-2 (Taipale et al., 2020), pinpoint outbreak areas, and enable local epidemiological interventions that maximize human health, while minimizing the extent of restrictive isolation measures.

Currently, most testing for active SARS-CoV-2 infection is performed using viral RNA extraction followed by RT-qPCR to amplify and detect several highly conserved regions of the SARS-CoV-2 genome. The global capacity for testing using this approach, however, has been limited in several ways. Initially, access and supply of reagents and instruments were limited considering the surge in demand. Second, this protocol requires a number of hands-on steps that must be performed by trained professionals, hampering its scalability, although automated systems do increase scale. Third, while several sequencing-based PCR approaches have been proposed (https://docs.google.com/document/d/1kP2w_uTMSep2UxTCOnUhh1TMCjWvHEY0sUUpkJHPYV4, https://www.notion.so/Octant-SwabSeq-Testing-9eb80e793d7e46348038aa80a5a901fd-639fd74b2ff14daf9a3b78bac1c738b1), their throughput is constrained by the availability of required devices like thermocyclers. Finally, and most critically, even as some of these bottlenecks have been reduced by automation and better supply chains, massive, repeated population testing is hampered by the need to collect samples in centralized settings, and to process each of them individually.

Here, we demonstrate LAMP-Seq, a novel protocol that allows for population-scale testing using massively parallel RT-LAMP (Nagamine et al., 2002; Notomi et al., 2000) by employing sample-specific barcodes. This approach requires only three heating steps for each individual sample, followed by pooled processing, parallelized deep sequencing, and well-established computational analysis. By using a simple thermal protocol for processing individual samples and pooling many samples prior to resource-intensive steps, the requirement for specialized reagents, equipment, and labor is greatly reduced relative to alternative protocols. Unique tracking of hundreds of millions of samples as well as asynchronous testing logistics, including at-home collection, can be achieved by employing a compressed barcode space. We describe the design of LAMP-Seq, validation on clinical specimens, and simulated barcoding strategies. We estimate that the cost per sample would be < 20 USD based on current list-prices of off-the-shelf products (excluding labor and instrument costs), with a potential for at least 10-fold cost reduction through scaled sourcing of three enzymes (RTx, Bst 2.0, Bst 3.0). Most importantly, this approach is predicted to be scalable to hundreds of thousands of samples per day per sequencing facility and could be deployed in developing countries.

## Results

We devised LAMP-Seq as an approach for population-scale testing for SARS-CoV-2 infection with the following overall steps (**Fig. 1A-C**): a barcoded RT-LAMP reaction is performed on an unpurified or lysed swab sample with primers specific for the SARS-CoV-2 genome, which is followed by large-scale pooling of samples, PCR amplification with additional barcoding, deep sequencing, and data analysis to identify positive individuals (**Fig. 1A, B**; see below for detailed protocol). RT-LAMP reactions have been demonstrated to be highly sensitive for sequence-specific viral nucleic acid detection (Lamb et al., 2020; Yang et al., 2020; Zhang et al., 2020), even from unpurified samples (Estrela et al., 2019). To establish a barcoded RT-LAMP reaction, we inserted barcode sequences into the forward inner primer (FIP), which enables generation of barcoded palindromic amplification products (**Fig. 1C**). When a small fraction of samples is expected to be positive during population scale testing, we can further limit the number of unique barcode primers needed for testing a large number of samples, by using a compressed barcode space (below).

**Figure 1.**
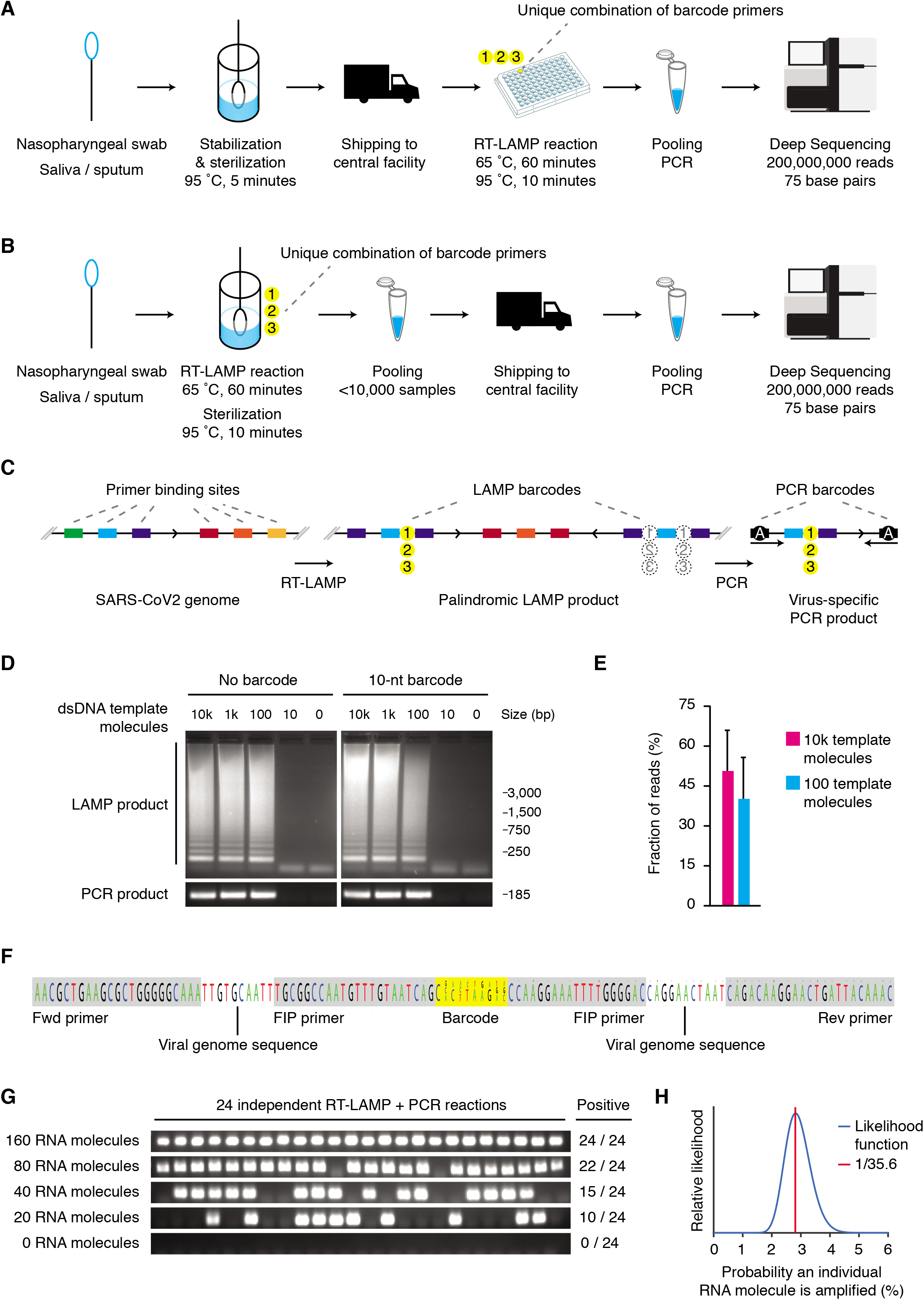
LAMP-Seq: A scalable deep-sequencing based approach for SARS-CoV-2 detection. (A) Schematic outline of a proposed scalable testing procedure involving remote lysis and inactivation of samples, and centralized barcoded RT-LAMP, pooling, and sequencing. (B) Schematic outline of a proposed scalable testing procedure involving remote barcoded RT-LAMP and sample pooling, and centralized sequencing. (C) Schematic of anticipated enzymatic reactions and reaction products. (D) Experimental validation of barcode insertion into FIP primers employed in LAMP reactions. All steps were performed as described in the Methods section, with the exception that plasmid DNA containing the SARS-CoV-2 N-gene (IDT) was used as template instead of a swab sample, no Bst 3.0 or Tris buffer were added, and the reaction was scaled down to a volume of 25 *μ*l. Samples were run on a 1% agarose gel and visualized using ethidium bromide. (E) Barcoded LAMP reactions templated with either 100 or 10,000 dsDNA molecules were combined after heat inactivation as described for D. Reactions were PCR amplified and sequenced on an Illumina MiSeq sequencer. Relative read counts are shown as mean and standard deviation from two experimental replicates. (F) RT-LAMP reactions with a combination of three barcoded FIP primers, but without Tris or Bst 3.0, were templated with synthetic RNA, and were sequenced using a MiSeq sequencer. Base frequencies are depicted by the size of each letter without applying any read filtering. Increasing phasing noise towards the 3’ end of the amplicon is likely caused by indels in primers. (G) Sensitivity measurement of RT-LAMP reactions as described for D templated with indicated numbers of synthetic RNA molecules. After PCR-amplification, the number of positive reactions was counted on a 1% agarose gel. (H) Likelihood function of the probability of detection for a single RNA molecule.

Specifically, we designed three barcoded primer sets based on validated RT-LAMP amplicons (**Table S1**, (Broughton et al., 2020; Lamb et al., 2020; Zhang et al., 2020)) perfectly matching 95.0% (amplicon A), 95.4% (amplicon B), and 96.8% (amplicon C) of 4,406 SARS-CoV-2 genomes available in the NCBI database (May 30th, 2020). 10-nt barcodes with GC content of 30%-70% and lacking homopolymer repeats of four or more nucleotides were inserted into the FIP primer. We ensured that barcodes are robust to sequencing errors by a minimum Levenshtein edit distance between any barcode pair sufficient to detect two insertion, deletion or substitution errors (**Table S1**).

Comparing barcoded LAMP reactions to non-barcoded controls using a dsDNA surrogate template for SARS-CoV-2, we confirmed that the presence of a prototypical 10-nt barcode within the FIP primer did not affect LAMP sensitivity, product amounts, or downstream PCR amplification (**Fig. 1D**). Templating two individually barcoded LAMP reactions that differ 100-fold in the amount of dsDNA template, combining them for PCR amplification, and sequencing the products resulted in read numbers within a two-fold range between the two samples (**Fig. 1E**), indicating that RT-LAMP saturation can effectively compress the dynamic range of input viral loads. This might be beneficial when analyzing many samples together on a sequencing run. Furthermore, we confirmed the expected sequence of barcoded RT-LAMP-PCR products by Illumina sequencing (**Fig. 1F**). In order to determine the molecular sensitivity of barcoded RT-LAMP reactions, we performed 24 reactions with differing numbers of template RNA molecules, and determined positive subsequent PCR reactions by gel electrophoresis (**Fig. 1G**). Using a constant per-molecule probability model of RNA detection, the maximum likelihood estimate for molecular detection efficiency is 1/35.5 per RNA molecule (**Fig. 1H**) which corresponds to an LoD-95 of 105 molecules. This is about an order of magnitude less sensitive than RT-qPCR (https://www.fda.gov/media/134922/download).

To validate LAMP-Seq, we tested 28 human samples side-by-side with a standard clinical diagnostic by RT-qPCR with a human subjects protocol approved by the ethics committee of the University Hospital Bonn. Two oropharyngeal samples were collected from each individual using two separate cotton swabs, which were anonymized using an individual ID. One swab was analyzed using a standard clinical diagnostics pipeline comprising rehydration, robotic RNA purification, and RT-qPCR (**Fig. 2A**, upper panel). The other swab was immediately inserted into a tube containing QuickExtract lysis buffer (Joung et al., 2020) (**Fig. 2A**, lower panel), processed and sequenced according to the LAMP-Seq protocol using individual PCR barcodes (**Methods**). The two methods were in complete agreement on both positives and negatives (**Fig. 2B, C**): All 12 individuals identified as SARS-CoV-2 RNA positive by RT-qPCR were also detected positive using LAMP-Seq employing a threshold of 10,000 reads; the remaining 16 individuals were identified as negative for viral RNA in agreement between both methods (**Fig. 2B, C**), with an average of 962 LAMP-Seq reads per negative sample, putatively arising from barcode swapping. Of note, N-gene-specific primers have been reported to be slightly less sensitive in RT-qPCR than primers for other targets (Corman et al., 2020). Unfiltered LAMP-Seq sequencing data confirmed the expected read structure, comprising primer sequences, viral genome sequence, and a matching barcode in 67% of reads (**Fig. 2D**), while the majority of remainder reads bore single-nucleotide substitutions or truncations relative to the expected amplicon sequence.

**Figure 2.**
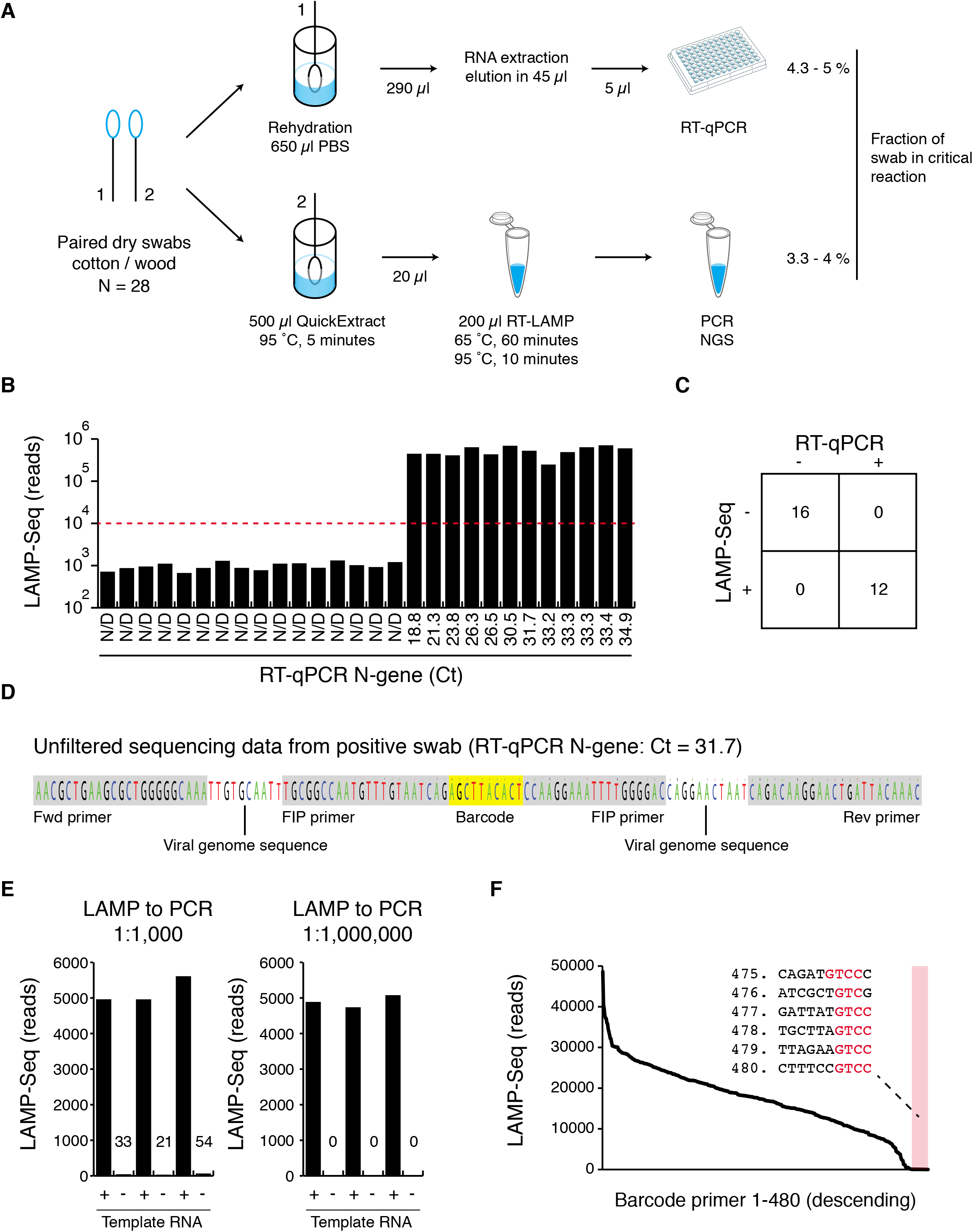
Clinical validation and optimization of LAMP-Seq. (A) Outline of the protocol employed for validating LAMP-Seq (lower workflow) against an established clinical RT-qPCR pipeline (upper workflow). (B) LAMP-Seq read numbers obtained per sample in comparison to RT-qPCR Ct values. The red dashed line indicates a threshold of 10,000 reads. (C) Summary statistics of validation experiments detailed in A, B. (D) NextSeq data obtained from a SARS-CoV-2-positive swab sample employing LAMP-Seq. Base frequencies are depicted by the size of each letter without applying any read filtering. (E) Quantitative assessment of barcode swapping during LAMP-Seq and its dependence on pre-dilution of pooled RT-LAMP reactions before PCR (left panel, 1,000-fold, right panel, 1,000,000-fold). LAMP-Seq was performed as described in the Methods section, with the exception that synthetic RNA was used as template instead of a swab sample, no Bst 3.0 or Tris buffer were added, and the reactions were scaled down to a volume of 25 μl. Numbers in the plot indicate read numbers for non-templated negative control reactions. (F) Empirical performance assessment of 480 random LAMP-Seq barcode primers. 480 barcoded FIP primers were mixed at equimolar concentration and were used as a pool in four replicate LAMP-Seq reactions templated by synthetic RNA. Raw sequencing data were analyzed using LAMP-Seq-Inspector v1.0 (http://manuscript.lamp-seq.org/Inspector.htm). Read counts are shown for barcodes in descending order, with the worst-performing 5 % of all barcode sequences highlighted in light red and the worst-performing barcode sequences listed.

Effective SARS-CoV-2 virus inactivation in QuickExtract lysis buffer was confirmed both after 30 minutes of incubation at 65 °C and after 10 minutes at 95 °C, resulting in at least a 3.9E4-fold reduction in viral infectivity (Table 1), whereas residual SARS-CoV-2 infectivity was retained following 30 minutes incubation at 22 °C. To further investigate the inactivation efficiency of the lysis buffer, a high dose of VSV virus was used to demonstrate at least a 1E7-fold reduction in viral infectivity (Table 1).

**Table 1.** Viral inactivation in QuickExtract lysis buffer. CPE, cytopathic effect.

| Incubation parameters |  | Viral inactivation |  |  |  |  |
| --- | --- | --- | --- | --- | --- | --- |
| Temperature | Time | Virus | Viral inoculation (log 10) | Residual viable virus (CPE) | Limit of detection (log 10) | Validated log 10 reduction |
| 65 °C | 20 min. | VSV | 9 | no | 2 | >7 |
| 22 °C | 30 min. | SARS-CoV-2 | 6 | yes | 2 |  |
| 65 °C | 30 min. |  | 6 | no | 2 | >4.6 |
| 95 °C | 10 min. |  | 6 | no | 2 | >4.6 |

We next optimized LAMP-Seq to allow successful pooling of barcoded RT-LAMP reactions, which is essential for scaling up LAMP-Seq, focusing on minimizing levels of barcode swapping, and on ensuring a sufficient number of individually validated barcodes. When we pooled six barcoded RT-LAMP reactions, of which three were templated with RNA, and performed PCR and sequencing, we observed moderate levels of barcode swapping (**Fig. 2E**, left panel). We hypothesized that barcode primers being transferred into the PCR reaction may lead to amplification and re-barcoding of amplicons. We eliminated detectable barcode swapping by diluting pooled RT-LAMP reactions one-million-fold in the PCR reaction, which (**Fig. 2E**, right panel). Next, pooling 480 barcoded FIP primers, performing RT-LAMP reactions in four replicates, and sequencing the barcode distribution in resulting products revealed that ~5% of barcode sequences perform poorly or even fail to engage in LAMP-Seq (**Fig. 2F**). Investigating potential sequence determinants that could guide optimized barcode design, we observed that the least efficient barcode primers displayed a marked enrichment for a GTCC motif or truncations thereof, especially towards the 3’ end of the barcode (**Fig. 2F**, inset). As this motif is the reverse complement of the 3’ end of the FIP primer, we hypothesize it could sequester the 3’ end by forming an intramolecular structure, thus inhibiting elongation of the primer, and should be avoided. Following this rule, we designed >6,000 barcoded FIP and BIP primers as well as provide 240 validated barcoded FIP primers for application of LAMP-Seq (**Table S1**).

A high-output Illumina 75-cycle NextSeq run can routinely generate 200 million sequencing reads in 14 hours, which we predict is sufficient for 100,000 samples per run, even accounting for library skewing due to differences in viral loads (for modeling see **Supplementary Note 1 and 2**). Barcoding 100,000 samples could be achieved by a naïve approach, where each sample is contacted with a unique barcode primer (**Fig. 3A**, left). However, as synthesis, validation, and robotic handling of large numbers of barcode primers is challenging, we explored a compressed barcode space, where every sample would be assigned a unique combination of more than one barcode (**Fig. 3A**, right). For this scenario, we conservatively assume that 1% of synthesized barcode primers systematically fail to work, even after removing barcode primers that contain homology to GTCC (Δ_synth_ = 0.01), and that 5% of all sample-specific barcodes are not detected due to varying sequencing depth (Δ_stoch_ = 0.05; independent of dropout due to low viral load). For automated assembly of testing reactions with unique barcode combinations, we anticipate that m = 1,000 barcode primers can be easily handled by available pipetting robots. Under these assumptions, we investigated for 100,000 samples what number of barcodes per sample (*k*), total number of barcode primers (*m*), and number of pools per run (*m_2_*) would minimize false-positive and false-negative rates of detection (**Fig. 3B-C**). Interpreting the compressed barcoding problem as a modified Bloom filter (**Supplementary Note 1**), we predict that when using *k* = 5 barcodes per sample, where *k’* = 3 barcodes are detected per sample, and splitting samples into *m2* = 96 pools per run, both the false-negative and false-positive rates of detection using a compressed barcode space will be less than 0.2% as long as the global frequency of positive samples is below 1.2% (**Fig. 3B**). Larger numbers of barcodes will further lower error rates and ensure performance in the face of higher global positive frequencies (**Fig. 3C**).

**Figure 3.**
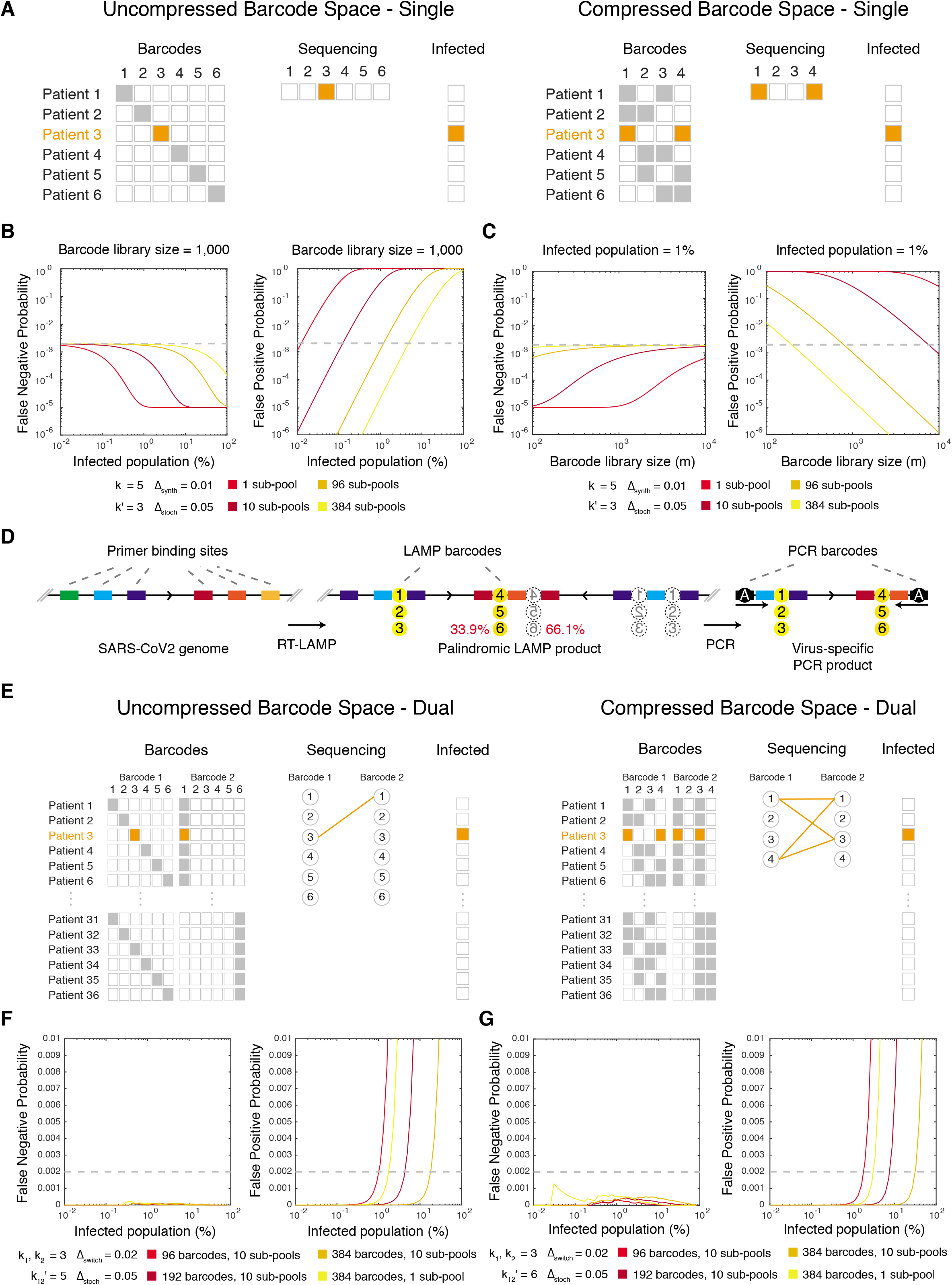
Modeling of compressed barcoding schemes for LAMP-Seq, enabling population-scale testing. (A) Schematic illustration of an uncompressed and a compressed single barcode scheme. (B-C) Calculated False Positive Probability and False Negative Probability depending on the global positive frequency of samples (A) and m = the complexity of the barcode library (B) for various numbers of pools per run, utilizing a compressed single barcode space accounting for barcode loss. Dashed grey lines indicate a probability threshold of 0.2%. (D) Schematic of anticipated enzymatic reactions and reaction products for dual barcoding. The depicted positioning of barcodes 4-6 in LAMP products is not the only conformation, with an alternative orientation indicated by dotted circles. (E) Schematic illustration of an uncompressed and a compressed dual barcode scheme. (F-G) Numerical simulation of False Positive Probability and False Negative Probability depending on the global positive frequency of samples, m_1_ = the number of FIP barcodes, m_2_ = the number of BIP barcodes, the number of pools per run, and k_12_’=the number of required barcode pairs per positive sample, over 100 iterations. Dashed grey lines indicate a probability threshold of 0.2%.

As some barcoded FIP primers fail in the RT-LAMP reaction, it may be advantageous to reduce the number of barcode primers that need to be validated. One way to achieve this is with a dual barcoding scheme, where both the FIP and BIP primers are barcoded (**Fig. 3D**). Using Tn5-mediated tagmentation and sequencing of RT-LAMP products (Thi et al., 2020), we experimentally quantified the formation of RT-LAMP products with the FIP/BIP barcode insertion sites facing each other to occur with a frequency of 33.1%, which suffices for PCR amplification of barcode pairs (**Fig. 3D**, red numbers). Without compression, 100,000 patient samples could be uniquely barcoded using 100 FIP primers, 100 BIP primers, and 10 pools per run. With compression, over 20 billion samples can each be assigned a unique combination of barcodes using a combination of 3 FIP primers and a combination of 3 BIP primers per patient sample from a pool of 96 barcoded FIP and 96 barcoded BIP primers (**Fig. 3E**).

This dual barcoding scheme would eliminate errors due to systematic barcode failure, but introduces the possibility of template switching errors. To explore parameters for this scheme, we assume that 5% of all sample-specific barcode pairs are not detected due to varying sequencing depth (Δ_stoch_ = 0.05; this is independent of dropout due to low viral load) and that template switching occurs 2% of the time (Δ_switch_ = 0.02). Under these assumptions, numerical simulations of this dual barcoding scheme (**Fig. 3F,G**, **Supplementary Note 2**) suggests that both the false-negative and false-positive rates of detection will be less than 0.2% as long as the global frequency of positive samples is below 1.6% when using a set of 96 FIP barcodes (m_1_, = 96), 96 BIP barcodes (m_2_ = 96), with 3 of each barcode per patient sample (*k*_1_, *k*_2_ = 3), requiring 6 out of 9 barcode pairs to be detected for a positive sample (*k*_12_’ = 6), and 10 pools per run (*m_3_* = 10) (**Fig. 3G**). Increasing the number of barcoded FIP and BIP primers to 192 or 384 each and changing the required threshold to 5 out of 9 barcode pairs detected for a positive sample (*k*_12_’ = 5) lowers the error rates and allows for higher global frequencies of positive samples (**Fig. 3F**). We emphasize that the simulated compressed barcoding schemes have not been experimentally validated yet.

## Discussion

LAMP and RT-LAMP (Nagamine et al., 2002; Notomi et al., 2000) have been previously established for use as highly sensitive methods for pathogen detection from unpurified human samples with detection limits below 100 nucleic acid molecules (Mori and Notomi, 2009). Although colorimetric or turbidimetric readouts of LAMP reactions can suffer from false positive results (Estrela et al., 2019), here we demonstrate that a sequencing-based readout provides maximum specificity by detecting only correct fusions of barcode sequences with two stretches of viral sequence. In addition, we show that this novel multiplexing-LAMP strategy can be made robust against barcode cross-contamination originating from template switching events or primer contamination at the PCR stage, as two template switching events would be required in order to create a sequencing-compatible amplicon.

We developed and optimized a barcoded RT-LAMP protocol (LAMP-Seq) and successfully validated it on 28 human swab samples. The current protocol does not require RNA purification or individual processing steps except using approximately one pipette tip per sample, which can be automated through using matrix-format tubes at the stage of swab lysis. Of note, larger numbers of patient samples need to be tested before proposing deployment of LAMP-Seq for population screening. Larger sample sizes will also allow exploration of the possibility of rare inhibitory compounds in some unpurified human samples, potentially resulting from food intake, hygiene interventions, or the oral microbiome. Apart from further validation studies, compatibility of the current LAMP-Seq protocol with other types of human samples (saliva, sputum, anterior nasal (AN) swabs, mid-nasal swabs, fecal samples) should be explored rapidly to identify the most scalable solution for unsupervised at-home sample collection, which would be attractive if the safety can be guaranteed during shipping of inactivated samples. For deployment, LAMP-Seq also has to be equipped with a positive control amplicon to ensure efficient RT-LAMP processing of each individual sample, which could run in the same RT-LAMP reaction or in a separate reaction, allowing independent saturation of both amplicons. Of note, the compressed barcoding schemes would require the positive control template to bear an additional heterogeneous sequence portion.

A major advantage of LAMP-Seq is that barcoding is performed early in the protocol using a simple heating device (like an oven), whereas downstream processing of sequencing libraries is done on large pools of samples. To enable pooled processing, we show that multiple barcode sequences can be inserted into the forward inner primer (FIP) and / or backward inner primer (BIP) primer used during an RT-LAMP reaction, as long as a simple sequence motif is avoided in all barcode sequences. As all barcodes have to be experimentally validated for diagnostic use, we propose and mathematically simulate a dual-indexing scheme that would allow uniquely barcoding more than 100,000 samples per run while only requiring 192 validated barcoded FIP/BIP primers.

A potential limitation of the presented approach is that skewing of sample representation at the pooling stage may affect testing sensitivity. Although the LAMP reaction saturates in positive samples largely independent of template concentrations (**Fig. 1E**), thus equalizing the representation across positive samples in an advantageous manner, the reaction might also add random skewing to pooled samples when scaling to hundreds of thousands of samples; however, preliminary modeling suggests that pooling 100,000 samples per NextSeq run offers robust detection (**Supplementary Note 1 and 2**).

LAMP-Seq requires low amounts of consumables with the exception of three proprietary enzymes and buffer compositions; however, these enzymes could be mass-produced using *E. coli* or replaced by open-source alternatives. The established LAMP-Seq protocol used cotton-wood-swabs that are available in mass quantities for < 5 ct. each. The synthesis cost of the barcode primer library is low overall (5,000 USD total for 960 barcodes, < 10 ct. per sample), leaving point-of-test infrastructure, logistics, and robotics as putative cost driving items. Once successfully established, however, this infrastructure could rapidly counter future waves of viral spread or pandemic outbreaks. Of note, LAMP-Seq could uniquely allow multiplexing multiple targets (of different viruses) to enable scalable differential diagnostics.

Broadly similar approaches of barcoded isothermal amplification methods have been independently suggested by other researchers (https://hms.harvard.edu/news/soup-nuts; Thi et al.; 2020; Palmieri et al.; Wu et al., 2020). To facilitate open communication, we have set up a forum on www.LAMP-Seq.org. We welcome collaboration from any interested parties so we can join together and rapidly develop a solution to advance the fight against the coronavirus pandemic.

## Materials and Methods

### LAMP-Seq testing for SARS-CoV-2 N-gene

1. A freshly inoculated cotton dry swab (nerbe plus GmbH, 09-819-5000) is inserted into 500 μl of QuickExtract (Lucigen, QE09050) supplemented with 2 ng/μl RNase-free plasmid DNA (pX330, Addgene #42230) in a 15 ml Falcon tube, stored on ice for transport, incubated for at least 10 minutes at room temperature, and heated to 95 °C for 5 minutes.
2. A barcoded RT-LAMP reaction is performed, containing the following components:

a. 100 *μ*l 2x LAMP master mix (NEB, E1700L),
b. 60 *μ*l 1 M Tris-HCl pH 8.6,
c. 2 *μ*l RNase-free plasmid DNA (pX330, Addgene #42230, 100 ng/μl),
d. 20 *μ*l swab lysate from step 1,
e. 5 *μ*l Bst 3.0 (NEB, M0374L, 8,000 units/ml),
f. 1.6 μM total of a unique set of one to five barcoded C-FIP primers (TGCGGCCAATGTTTGTAATCAGNNNNNNNNNNCCAAGGAAATTTTGGGG AC), where Ns denote a specific barcode sequence,
g. 1.6 μM C-BIP primer (CGCATTGGCATGGAAGTCACTTTGATGGCACCTGTGTAG),
h. 0.2 μM C-F3 primer (AACACAAGCTTTCGGCAG),
i. 0.2 μM C-B3 primer (GAAATTTGGATCTTTGTCATCC),
j. 0.4 μM C-LF primer (TTCCTTGTCTGATTAGTTC),
k. 0.4 μM C-LB primer (ACCTTCGGGAACGTGGTT),
l. water to a total volume of 200 μl.
3. Optionally, the RT-LAMP reaction is split into eight reactions.
4. The RT-LAMP reaction is heated to 65 °C for 1 hour, and to 95 °C for 10 minutes.
5. Up to 100,000 reactions are pooled in batches of 1,000 to 10,000 samples per batch.
6. The pool is diluted 1:100,000 in water.
7. For each pool, a 20-cycle 50 *μ*l PCR reaction is performed:

a. 25 *μ*l NEBNext 2x Master Mix (NEB),
b. 0.5 *μ*M PCR-C-fwd primer (ACACTCTTTCCCTACACGACGCTCTTCCGATCTAACGCTGAAGCGCTGGGGGCAAA),
c. 0.5 *μ*M PCR-C-rev primer (TGACTGGAGTTCAGACGTGTGCTCTTCCGATCTGTTTGTAATCAGTTCCTT GTCTG),
d. 5 *μ*l of diluted RT-LAMP reactions from step 5,
e. water.
8. For each pool, a secondary 12-cycle 50 μl PCR reaction is performeds with:

a. 25 μl NEBNext 2x Master Mix (NEB),
b. 0.5 μM pool-specific fwd barcoding primer (AATGATACGGCGACCACCGAGATCTACACNNNNNNNNNNACACTCTTTCCCTACACGACGCT), where Ns denote a specific barcode sequence,
c. 0.5 μM pool-specific rev barcoding primer (CAAGCAGAAGACGGCATACGAGATNNNNNNNNNNGTGACTGGAGTTCAGACGTGTGCT), where Ns denote a specific barcode sequence,
d. 5 μl of previous PCR reaction,
e. water.
9. The PCR products are pooled on ice, purified using a silica spin column (Qiagen), quantified using a NanoDrop photospectrometer (Thermo) or Qubit (Thermo), and sequenced on an Illumina NextSeq sequencer or similar device (A MiSeq sequencer can be used for method testing, or when screening smaller numbers of samples).
10. Using the *LAMP-Seq-lnspector* software (http://manuscript.lamp-seq.org/Inspector.htm), barcodes co-occurring with the correct viral genome sequence excluding sequence portions covered by primers are determined. This analysis can also be performed using a “kallisto | bustools” workflow (Booeshaghi et al., 2020).
11. Positive samples are determined using a database of barcode combinations assigned to sample IDs, requiring either one (single barcoding scenario) or at least three out of five sample barcodes (compressed barcode space) being positive.

### Clinical RT-qPCR pipeline

Swabs were rehydrated in 650 *μ*l PBS. Viral RNA was extracted using the chemagic™ Prime Viral DNA/RNA 300 Kit (PerkinElmer) on a Chemagic Prime 8 system (PerkinElmer). 290 *μ*l viral sample were mixed with 10 *μ*l extraction control sample and 300 *μ*l lysis buffer. Extraction was performed according to the manufacturers protocol and viral RNA was eluted in 45 *μ*l elution buffer for subsequent analysis. Detection of viral RNA using one-step real-time reversetranscription PCR was performed according to (Corman et al., 2020) with iTaq Universal Probes One-Step Kit (Biorad) using primers and probes against the N-gene (N_Sarbeco_F: CACATTGGCACCCGCAATC, N_Sarbeco_R: GAGGAACGAGAAGAGGCTTG, N_Sarbeco_P: FAM-ACTTCCTCAAGGAACAACATTGCCA-BBQ, TIB MolBiol). Spike-in RNA of the bacteriophage MS2 served as an interal control and was detected with Luna^®^ Universal Probe One-Step RT-qPCR Kit (New England Biolabs) using corresponding primers and probe (MS2_F: TGCTCGCGGATACCCG, Ms2_R: AACTTGCGTTCTCGAGCGAT, MS2_P: YAK-ACCTCGGGTTTCCGTCTTGCTCGT — BBQ, TIB MolBiol). The reaction for the internal control was performed using dual detection of FAM and YAK/VIC in a Lightcycler 480 (Roche), the detection of the N-gene was done in a QuantStudio5 cycler (Thermo Fisher).

### Viruses and cells

SARS-CoV-2 strain MUC-IMB-1 was isolated and kindly supplied by Rosina Ehmann and Gerhard Dobler (Bundeswehr institute of microbiology, Munich, Germany). The virus was propagated and titrated on VERO-E6 cells (ATCC CRL-1586). All handling and working with SARS-CoV-2 was conducted in a BSL-3 facility in accordance with the biosafety guidelines of the IIBR. Vesicular stomatitis virus (VSV) serotype Indiana, kindly provided by Eran Bacharach (Tel-Aviv University, Israel), was propagated and titrated on Vero cells (ATCC CCL-81). Handling and working with VSV was conducted in a BSL-2 facility in accordance with the biosafety guidelines of the IIBR.

### Lysis buffer inactivation assay

Quick extract DNA extraction solution (Lucigen) was tested in accordance with the manufacturer’s suggested buffer to sample ratio. Universal Transfer Medium (UTM, Copan) aliquots were inoculated with either 5E6 pfu/ml SARS-CoV-2 or 2E9 pfu/ml VSV viruses and were incubated at 22 °C, 65 °C, or 95 °C for 10 to 30 minutes. Positive and negative control samples included UTM inoculated with viable virus without lysis buffer and UTM with Lysis buffer without virus, respectively. The limit of detection was defined as the first serial dilution of negative control that did not cause CPE by itself (represented in log scale). Briefly, VERO-E6 (for SARS-CoV-2) or VERO cells (for VSV) were cultured in DMEM supplemented with 10% FBS, MEM non-essential amino acids, 2 mM L-Glutamine, 100 U/ml penicillin, 0.1 mg/ml streptomycin, and 12.5 U/ml Nystatin (Biological Industries, Israel). Monolayers (2.5E5 cells per well in 24-well plates) were washed once with MEM Eagles medium without FBS, and infected with 200 *μ*l of ten-fold serial dilutions of the samples. After one hour of incubation the wells were overlaid with 1 ml of MEM medium containing 2% fetal calf serum (FCS), MEM non-essential amino acids, 2 mM L-Glutamine, 100 U/ml penicillin, 0.1 mg/ml streptomycin, 12.5 U/ml Nystatin, and 0.15% Sodium Bicarbonate (Biological Industries, Israel). The cells were then incubated at 37 °C, 5% CO_2_ for five days (SARS-CoV-2) or one day (VSV). CPE was determined by counter-staining with crystal violet solution.

### Code and Data Availability

The LAMP-Seq Inspector tool for processing raw LAMP-Seq data is available at: http://manuscript.lamp-seq.org/Inspector.htm. Python scripts for designing the error-correcting barcodes are available at: https://github.com/feldman4/dna-barcodes. Jupyter Notebooks for numerical simulations and MATLAB scripts for figure generation are available at: https://github.com/dbli2000/SARS-CoV2-Bloom-Filter. Example LAMP-Seq data is available on www.LAMP-Seq.org.

## Acknowledgements

Foremost, we thank all participating individuals who enabled this research by donating swab samples. We thank Rhiannon Macrae for help with preparing the manuscript, and Michael Knop, George Smith, Gunther Hartmann, Phillip Buckhaults, Lior Pachter, Yuval Dor, Sophie Strobel, Samantha Laber, Amy Guo, André Heimbach, Hendrik Streeck, Eric Zhang, Vincent Huang, Daniel Liu, Abhishek Vijayakumar, Sebastian Virreira Winter, Johannes Schmid-Burgk, Stefan Frank, and José Miguel Zapata Rolón for helpful discussions. We thank Lara Hochfeld, Nina Ishorst, Eva Beins, and Peter Teßmann for help performing the RT-qPCR, and Daniel Hinze for plasmid preparation. M.H. and M.M.N. were supported by the Deutsche Forschungsgemeinschaft (DFG, German Research Foundation) under Germany’s Excellence Strategy - EXC2151 - 390873048. K.L. and M.M.N. are members of the German COVID-19 OMICS initiative (DeCOI, https://decoi.eu/). AR and FZ are Investigators of the Howard Hughes Medical Institute. Work was supported by the Klarman Incubator (AR).

## References

Booeshaghi, A.S., Lubock, N.B., Cooper, A.R., Simpkins, S.W., Bloom, J.S., Gehring, J., Luebbert, L., Kosuri, S., and Pachter, L. (2020). Fast and accurate diagnostics from highly multiplexed sequencing assays. medRxiv 2020.05.13.20100131.

Broughton, J.P., Deng, X., Yu, G., Fasching, C.L., Singh, J., Streithorst, J., Granados, A., Sotomayor-Gonzalez, A., Zorn, K., Gopez, A., et al. (2020). Rapid Detection of 2019 Novel Coronavirus SARS-CoV-2 Using a CRISPR-based DETECTR Lateral Flow Assay. 1–27.

Chinazzi, M., Davis, J.T., Ajelli, M., Gioannini, C., Litvinova, M., Merler, S., Pastore y Piontti, A., Mu, K., Rossi, L., Sun, K., et al. (2020). The effect of travel restrictions on the spread of the 2019 novel coronavirus (COVID-19) outbreak. Science eaba9757–12.

Corman, V.M., Landt, O., Kaiser, M., Molenkamp, R., Meijer, A., Chu, D.K., Bleicker, T., Brünink, S., Schneider, J., Schmidt, M.L., et al. (2020). Detection of 2019 novel coronavirus (2019-nCoV) by real-time RT-PCR. Euro Surveill. 25, 2431.

Dong, E., Du, H., and Gardner, L. (2020). An interactive web-based dashboard to track COVID-19 in real time. Lancet Infect Dis 20, 533–534.

Estrela, P.F.N., de Melo Mendes, G., de Oliveira, K.G., Bailão, A.M., de Almeida Soares, C.M., Assunção, N.A., and Duarte, G.R.M. (2019). Ten-minute direct detection of Zika virus in serum samples by RT-LAMP. Journal of Virological Methods 271, 113675.

Joung, J., Ladha, A., Saito, M., Segel, M., Bruneau, R., Huang, M.-L.W., Kim, N.-G., Yu, X., Li, J., Walker, B.D., et al. (2020). Point-of-care testing for COVID-19 using SHERLOCK diagnostics. medRxiv 2020.05.04.20091231.

Lamb, L.E., Bartolone, S.N., Ward, E., and Chancellor, M.B. (2020). Rapid Detection of Novel Coronavirus (COVID-19) by Reverse Transcription-Loop-Mediated Isothermal Amplification. 1–17.

Li, R., Pei, S., Chen, B., Song, Y., Zhang, T., Yang, W., and Shaman, J. (2020). Substantial undocumented infection facilitates the rapid dissemination of novel coronavirus (SARS-CoV2). Science 6, eabb3221–eabb3229.

Mori, Y., and Notomi, T. (2009). Loop-mediated isothermal amplification (LAMP): a rapid, accurate, and cost-effective diagnostic method for infectious diseases. Journal of Infection and Chemotherapy 15, 62–69.

Nagamine, K., Hase, T., and Notomi, T. (2002). Accelerated reaction by loop-mediated isothermal amplification using loop primers. Molecular and Cellular Probes 16, 223–229.

Notomi, T., Okayama, H., Masubuchi, H., Yonekawa, T., Watanabe, K., Amino, N., and Hase, T. (2000). Loop-mediated isothermal amplification of DNA. Nucl. Acids Res. 28, E63–E63.

Palmieri, D., Siddiqui, J.K., Gardner, A., Fishel, R., bioRxiv, W.M., 2020 REMBRANDT: A high-throughput barcoded sequencing approach for COVID-19 screening. Biorxiv.org

Taipale, J., Romer, P., and Linnarsson, S. (2020). Population-scale testing can suppress the spread of COVID-19. medRxiv 2020.04.27.20078329.

Thi, V.L.D., Herbst, K., Boerner, K., Meurer, M., Kremer, L.P.M., Kirrmaier, D., Freistaedter, A., Papagiannidis, D., Galmozzi, C., Klein, S., et al. (2020). Screening for SARS-CoV-2 infections with colorimetric RT-LAMP and LAMP sequencing. medRxiv 2020.05.05.20092288.

Wu, Q., Suo, C., Brown, T., Wang, T., Teichmann, S.A., and Bassett, A.R. (2020). INSIGHT: a scalable isothermal NASBA-based platform for COVID-19 diagnosis. bioRxiv 2020.06.01.127019.

Yang, W., Dang, X., Wang, Q., Xu, M., Zhao, Q., Zhou, Y., Zhao, H., Wang, L., Xu, Y., Wang, J., et al. (2020). Rapid Detection of SARS-CoV-2 Using Reverse transcription RT-LAMP method. 1–25.

Yu, Y., Liu, Y.-R., Luo, F.-M., Tu, W.-W., Zhan, D.-C., Yu, G., and Zhou, Z.-H. (2020). COVID-19 Asymptomatic Infection Estimation. medRxiv 2020.04.19.20068072.

Zhang, Y., Odiwuor, N., Xiong, J., Sun, L., Nyaruaba, R.O., Wei, H., and Tanner, N.A. (2020). Rapid Molecular Detection of SARS-CoV-2 (COVID-19) Virus RNA Using Colorimetric LAMP. 1–14.

