## Supplementary Note 1 for "LAMP-Seq: Population-Scale COVID-19 Diagnostics Using Combinatorial Barcoding"

#### LAMP-Seq: Population-scale COVID-19 Diagnostics Using Combinatorial Barcoding - Theory

We want to develop a method to scalably detect SARS-CoV-2-infected patients. Our goal here is to model and characterize error rates of various approaches for doing so.

For this supplementary note, we assume that there are two stages of barcoding available: patient samples will be individually barcoded at the RT-LAMP stage with a first set of barcodes (barcode 1), and then groups of patient samples will be barcoded with a second set of orthogonal barcodes (barcode 2) in the process of preparing samples for Illumina sequencing. After sequencing, a given barcode pair can be called as positive or negative for SARS-CoV-2 viral RNA. Let there be $m$ total unique barcode 1s and $m_2$ total unique barcode 2s.

Suppose that we want to test $N$ total patient samples, which will be done in batches of size $b$ with $k$ pre-assigned barcode 1s per patient sample. Further suppose that a fraction $p$ of the total population is positive for SARS-CoV-2. Then, $\approx n = pb$ samples in each batch will be positive. There are two types of general approaches considered below, one where $k = 1$ and one where $k > 1$ (corresponding to the single index barcode schemes in the main text).

To compute error rates for any approach, we must have a model for how errors arise. We assume that errors from barcode loss and sample skewing dominate, modeling barcode loss first and then sample skewing.

#### 1 Modeling Barcode Loss

We assume that barcode loss occurs only for barcode 1s, and that there are two types of errors. This is because there are many copies of RT-LAMP product for a positive sample before addition of barcode 2, and because barcode 2 failure is easy to detect and correct due to the small number of barcode 2s used in practice. We will also assume that no negative barcodes will show up as positive, as template switching with this barcoding scheme is unlikely.

The first type is an error where a given barcode never functions properly, which will happen with probability $\Delta_{synth}$. This may be because it was not synthesized properly or because it impedes amplification. In practice, some initial barcoded FIP primers tested did not work with RT-LAMP. Barcodes were designed with at least one-bit error correction, so small synthesis errors are likely negligible. Overall, after filtering, we expect this to be low ($\approx 0.01$).

The second type is an error in which a given barcode may not be picked up for a particular positive sample with probability $\Delta_{stoch}$, perhaps due to dilution effects or liquid handling errors. We expect that this probability will vary with the number of people infected in a batch $n$, so it should also vary with $p$ and $b$. This probability has not been well characterized.

#### 2 Scenario 1: $k = 1$

If $b < m \cdot m_2$, then every sample in a batch can be assigned a unique barcode. It is challenging to synthesize enough barcodes for $N < m \cdot m_2$, which would enable a unique barcode for every sample from the total population, so batches would have to be defined in some way. This suggests that this scenario would likely require synchronous testing. Every patient sample in a batch would then have a unique barcode 1 - barcode 2 pair.

Since each patient has a unique barcode pair, there are no false positives. However, with barcode loss, there may be false negatives. Each patient sample would have a false negative probability of $\Delta_{synth} + \Delta_{stoch} (1 - \Delta_{synth})$.

#### 3 Scenario 2: $k > 1$

With a liquid handler, it may be possible to give each patient sample $k$ different barcode 1s, with $k > 1$, in a pre-assigned way. Then, if $N < \binom{m}{k}$, every patient sample for the entire population could have a unique combination of barcode 1s. This does not mean that every barcode 1 would correspond to a unique patient sample, as this is only possible if $N < \frac{m}{k}$.

For this scenario, we imagine an asynchronous sample collection system where $b$ of these samples, as they come in, are split into $m_2$ sub-batches. Then, each sub-batch gets a unique barcode 2. If, after sequencing, $\geq k'$ out of the $k$ barcode pairs that correspond to a specific patient sample come up as positive, we would say that that patient sample is positive.

Since the barcode 1s no longer correspond to unique patient samples, there is a potential for false positives using this approach. False negatives are also still possible, as a positive sample could be called as negative if more than $k - k'$ of the $k$ corresponding barcode pairs are lost. Using a modified Bloom filter model (see Appendix A, (1)), we can compute the false positive probability ($FPP_{\Delta k', m_2}$) and false negative probability ($FNP_{\Delta k', m_2}$) for this approach as

$$FPP_{\Delta k', m_2} = \sum_{i=k'}^{k} \binom{k}{i} \left( \Delta_{synth} + (1 - \Delta_{synth}) \left( 1 - \frac{1 - \Delta_{stoch}}{m} \right)^{\frac{kn}{m_2}} \right)^{k-i}$$
$$\left( (1 - \Delta_{synth}) \left( 1 - \left( 1 - \frac{1 - \Delta_{stoch}}{m} \right)^{\frac{kn}{m_2}} \right) \right)^{i}$$

$$FNP_{\Delta k', m_2} = 1 - \sum_{i=k'}^{k} \binom{k}{i} \left( \Delta_{synth} + (1 - \Delta_{synth}) \Delta_{stoch} \left( 1 - \frac{1 - \Delta_{stoch}}{m} \right)^{k\left(\frac{n}{m_2} - 1\right)} \right)^{k-i}$$
$$\left( (1 - \Delta_{synth}) \left( 1 - \Delta_{stoch} \left( 1 - \frac{1 - \Delta_{stoch}}{m} \right)^{k\left(\frac{n}{m_2} - 1\right)} \right) \right)^{i}.$$

Note that these error rates are both dependent on the proportion of the population infected, $p$, since $n = pb$. As $p$ increases, the false negative probability will drop, as other positive patient samples can compensate for the barcode loss of a given sample. We can compute an upper bound on the false negative probability ($FNP_{\Delta k', m_2, max}$) that avoids this $p$ dependence as

$$FNP_{\Delta k', m_2, max} = 1 - \sum_{i=k'}^{k} \binom{k}{i} \left( \Delta_{synth} + (1 - \Delta_{synth}) \Delta_{stoch} \right)^{k-i} \left( (1 - \Delta_{synth}) (1 - \Delta_{stoch}) \right)^{i}.$$

Additionally, the barcode loss lowers the false positive probability in this scenario. Since $\Delta_{stoch}$ and $\Delta_{synth}$ are both not fully characterized, it may be worth considering the case where there is no barcode loss ($\Delta_{stoch} = \Delta_{synth} = 0$) for robustness. Then, the false positive probability, ($FPP_{\Delta k', m_2, max}$), which is still a function of $p$, can be given as

$$FPP_{\Delta k', m_2, max} = \sum_{i=k'}^{k} \binom{k}{i} \left( \left( 1 - \frac{1}{m} \right)^{\frac{kn}{m_2}} \right)^{k-i} \left( 1 - \left( 1 - \frac{1}{m} \right)^{\frac{kn}{m_2}} \right)^{i}.$$

Overall, a lower $k'$ for a fixed $k$ leads to a reduced false negative probability at the cost of an increased false positive probability. The effect of varying $k$ depends on the chosen $k'$ and on the proportion of the population infected $p$.

Analysis of scenario 2 using realistic numbers can be found in the main text. Some additional error plots with other values for $m$ are shown in Figures 1 and 2.

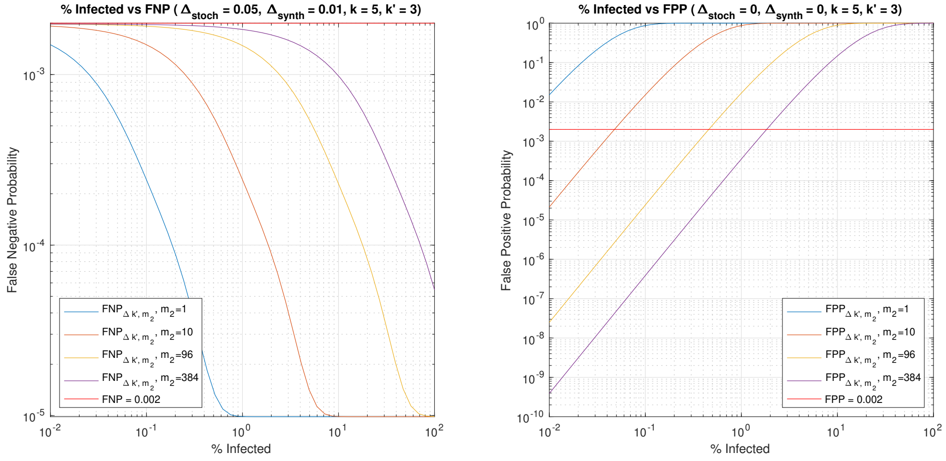


Figure 1: Error rates for various $m_2$ as the % infected varies for $m = 384$ and $b = 10^5$.

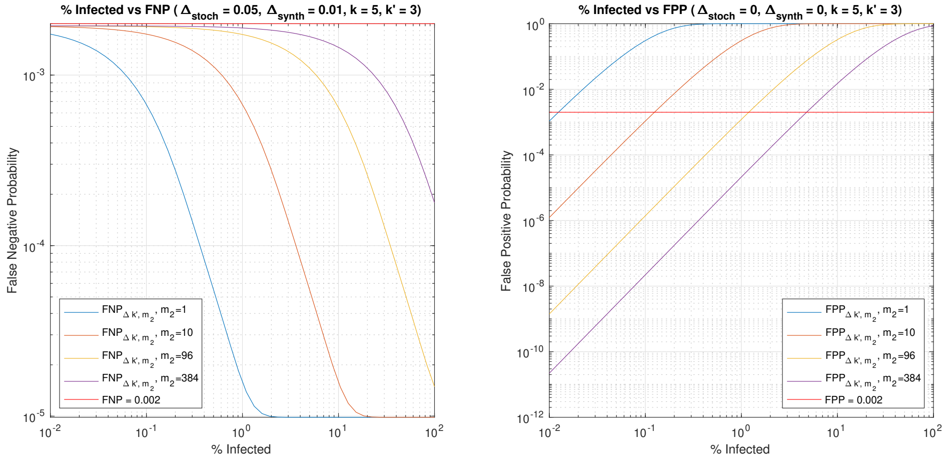


Figure 2: Error rates for various $m_2$ as the % infected varies for $m = 10^3$ and $b = 10^5$.

### 4 Modeling Sample Skewing

Sample skewing errors could also occur. Patient viral loads from nasopharyngeal swabs vary over many orders of magnitude across the course of infection (2). This variation could lead to over-representation of some positive samples, preventing detection of samples with lower viral abundance and giving rise to false negatives.

RT-LAMP and PCR are both nonlinear amplification methods, and how patient viral load variation propagates through both techniques remains to be experimentally determined. To initially model this using numerical simulation, we consider two possibilities.

One possibility is that the saturation of both RT-LAMP and PCR lead the distribution of barcoded molecules post-PCR to have less variation than the original viral load. We model this by drawing the molecules post-PCR for a barcode from a normal distribution with mean $10^4$ and standard deviation $10^3$, with the same number of molecules for each of the $k$ barcodes for a positive sample. This is referred to as "Saturated". Initial data collection suggests that LAMP-seq leads to saturation, so we assume this is the more accurate model.

Another possibility is that the RT-LAMP and PCR lead to the retention or exacerbation of the initial sample variation. We model this by drawing the molecules post-PCR for a barcode from a log-normal distribution with mean 4.5 and standard deviation 3, with the same number of molecules for each of the $k$ barcodes for a positive sample. This is referred to as "Amplified".

Overall, the modeled molecules per sample is not important, and may be orders of magnitude higher, as the relative abundance of various barcodes is what will determine false negatives.

Assuming that an Illumina NextSeq run generates 200 million total sequencing reads and that $\approx 90\%$ of the reads are usable, there are a total of $\approx$ 18 million reads per sub-batch. In the simulation, a given barcode was called as positive if the number of reads for the barcode (calculated by relative abundance multiplied by reads per sub-batch) was greater than or equal to a threshold of $t$ reads. Otherwise, it was called as a negative barcode.

Using the parameters for Scenario 1 and 2 ($k' = 3$ and $k = 5$) without barcode loss, we calculated the average FPPs and FNPs over 1000 iterations for various threshold values $t$, shown in Table 1. Under the "Saturated" model, sample skewing introduces minimal false negatives. With a threshold of 100 reads, the FNP is tolerable for both scenarios. For the "Amplified" model, sample skewing introduces more false negatives, with the threshold reads $t$ determining exactly how many. As the proportion of the population infected $p$ goes up, the FNP will increase.

| Threshold Reads $t$ | Average FNP | Average FPP | Model | Scenario |
|---|---|---|---|---|
| 100 | 0.0000 | 0.0000 | Saturated | 1 |
| 2 | 0.0087 | 0.0000 | Amplified | 1 |
| 10 | 0.0296 | 0.0000 | Amplified | 1 |
| 100 | 0.1227 | 0.0000 | Amplified | 1 |
| 100 | 0.0000 | 0.0010 | Saturated | 2 |
| 2 | 0.0289 | 0.0010 | Amplified | 2 |
| 10 | 0.0817 | 0.0009 | Amplified | 2 |
| 100 | 0.2567 | 0.0005 | Amplified | 2 |

Table 1: FPP/FNP for various $t$ with $b = 10^5, m = 10^4, m_2 = 10$, and $p = 0.01$.

However, it is unclear how accurate the "Saturated" model is. The propagation of sample variation through LAMP-seq needs to be experimentally determined, in order to refine this model and to guide parameter choice.

We assumed here that either sample skewing or barcode loss dominates. Combining both may produce a more realistic model. Simulations doing so are available in the Github repository.

### 5 Other Potential Approaches

The two scenarios presented here are not the only possible approaches with this barcode scheme, and others may have lower error rates. Some scenarios where either the barcode 1s or barcode 2s are not utilized are modeled in Appendix $A$ and some more complex scenarios with redundancy across barcode 2s are briefly explored in Appendix $B$.

### 6 Code Availability

The scripts used to simulate and plot the models described here are available at the Github repository `https://github.com/dbli2000/SARS-CoV2-Bloom-Filter`.

### References


[1] Burton H. Bloom. 1970. *Space/time trade-offs in hash coding with allowable errors.* Commun. ACM 13, 7 (July 1970), 422–426. DOI: https://doi.org/10.1145/362686.362692

[2] Wölfel, R. et al. *Virological assessment of hospitalized patients with COVID-2019.* Nature (2020). DOI: https://doi.org/10.1038/s41586-020-2196-x

### A Modeling as a Modified Bloom Filter

A useful first approximation for this problem is that of a Bloom filter. Using an idealized Bloom filter, we first calculate some theoretical limits and optima if we only use barcode 1s. We then modify the Bloom filter to incorporate barcode loss, as well as different criteria for calling a sample as positive, before adding in the barcode 2s to produce a final model. All derivations can be found in Appendix C.

#### A.1 Bloom Filters and Correspondence to the Problem

A Bloom filter is a data structure that can be used to test whether an element is a part of a set of cardinality $n$ (1). This test returns either "definitely not", or "possibly yes". A Bloom filter is implemented by taking a bit array of size $m$, initialized at 0. To add an element to the filter/set, $k$ hash functions are used to map the new element to $k$ bits in the array. These $k$ bits are then set to 1, if they are not already 1. To query whether a given element is in the set, compute the $k$ hashes again. If any of the corresponding $k$ bits in the array are 0, then the element is not in the set. If they are all 1, then the element may be in the set.

We will assume that there are no barcode 2s for now (i.e. $m_2 = 1$). Then, a Bloom filter can be generated for every batch of size $b$ we test. Here, elements are individual samples, and inclusion in the set of cardinality $\approx n = pb$ corresponds to being SARS-CoV-2 positive. The $m$ barcodes correspond to the $m$ bits in the bit array, where being set to 0/1 means it was detected as SARS-CoV-2 negative or SARS-CoV-2 positive respectively. Similarly, the outputs of the $k$ hash functions are equivalent to the $k$ barcodes assigned to a given sample.

#### A.2 Underlying Assumptions

The critical assumption of this model that is broken in real life is the idea that a barcode will always turn up positive if it is positive, implying no barcode loss. This is addressed and modeled in later sections, but it still useful to consider this initial idealized case for now.

Otherwise, the other assumptions mostly hold. Namely, the $k$ hash functions are supposed to be independent and produce a random uniform distribution across the keys (different barcodes). This can be achieved using a proper barcode assignment. Another assumption is that barcodes will not spontaneously turn positive, which should be the case under this barcoding scheme. Bits are assumed to be set to either 0 or 1, which is realistic if we binary threshold the reads for a given barcode.

#### A.3 Single Bloom Filter, No Barcode Loss

##### A.3.1 False positives/negatives

For this idealized case, if an element is in the set, all $k$ bits will be set to 1, so false negatives are not possible.

However, if, by chance, all $k$ corresponding bits in the filter for an element $e$ are set to 1 due to other elements that combined to map to the same $k$ bits, we will mistakenly claim that $e$ is in the set, even if it is not. So **false positives are possible**, analogous to hash collisions. This false positive probability ($FPP$) is

$$FPP = \left(1 - \left(1 - \frac{1}{m}\right)^{kn}\right)^k \approx \left(1 - e^{-\frac{kn}{m}}\right)^k .$$

This makes certain assumptions about the hash functions, but these are $\approx$ satisfied by the barcode assignment (see Appendix C).

##### A.3.2 Optimal choice of $k$

There is an optimal choice for $k$ that minimizes the false positive rate. This value is closely approximated by $k_{OPT} = \ln(2)\frac{m}{n}$ with a corresponding false positive probability (FPP) of $0.5^{k_{OPT}}$. Note that what matters is the ratio $m$ to $n$, so things can scale accordingly with various numbers of barcodes as long as the batch size to barcode number ratio remains the same for a given % infected.

If we choose $m = 10,000$ barcodes and test batches of size $b = 100,000$ at once, then we can plot the optimal choice of $k$ given the percent of the population infected, as well as the corresponding false positive probability (Figure 3). We can also generate a table showing the optimal $k$ given a certain percentage of the population infected (Table 2).

When the percentage infected is low, $k_{OPT}$ is high, and leads to a low FPP. As the percentage infected rises, $k_{OPT}$ will drop, due to barcode saturation, and the FPP rises. So, in early stages, high values for $k$ are optimal, and this shrinks across the course of the pandemic. However, adjusting $k$ as the pandemic evolves may not be realistic. High $k$ may also cause technical problems with the RT-LAMP reaction. Instead, we will focus on what happens if we pre-choose $k = 2, 3, 4$ or 5.

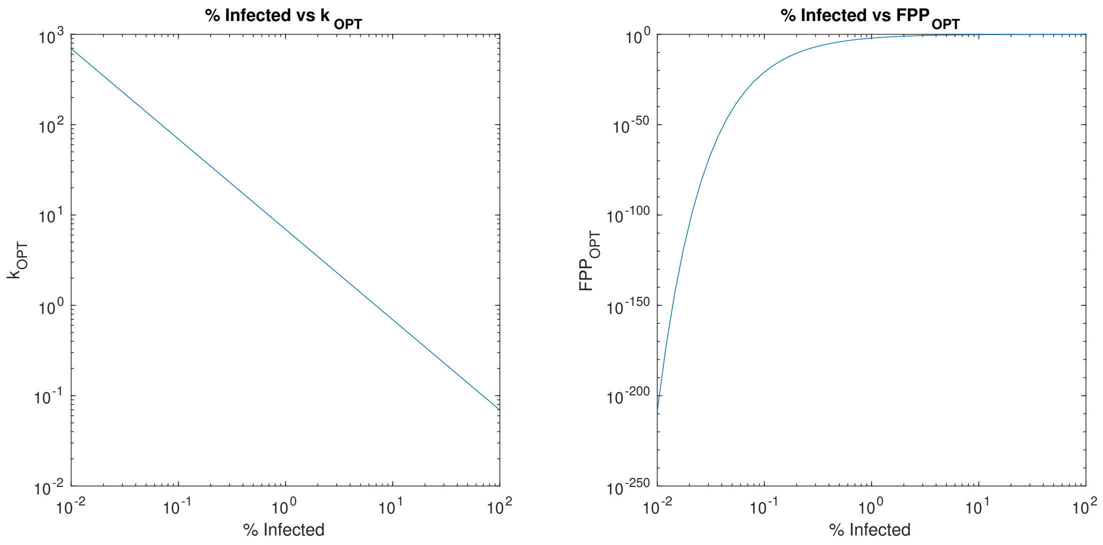


Figure 3: $k_{OPT}$ and $FPP_{OPT}$ as % Infected varies with $m = 10^4$ and $b = 10^5$

| $k_{OPT}$ | $n$ (Infected/Batch) | % Infected ($b = 10^4$) | % Infected ($b = 10^5$) | FPP |
|---|---|---|---|---|
| 138.63 | 50 | 0.5 | 0.05 | $1.85 \times 10^{-42}$ |
| 69.3 | 100 | 1 | 0.1 | $1.37 \times 10^{-21}$ |
| 34.66 | 200 | 2 | 0.2 | $3.68 \times 10^{-11}$ |
| 13.86 | 500 | 5 | 0.5 | $6.73 \times 10^{-5}$ |
| 6.93 | 1000 | 10 | 1 | 0.00820 |
| 5 | 1386 | 13.86 | 1.386 | 0.03125 |
| 4 | 1733 | 17.33 | 1.733 | 0.0625 |
| 3 | 2311 | 23.11 | 2.311 | 0.125 |
| 2 | 3466 | 34.66 | 3.466 | 0.250 |
| 1.38 | 5000 | 50.00 | 5 | 0.384 |

Table 2: $k_{OPT}$ and False Positive Probability (FPP) assuming $m = 10^4$ barcodes

### A.3.3 $k = 2, 3, 4$ or 5

When operating with $k = 2, 3, 4$ or 5, we are not very close to the current theoretical optimum. However, it may be the best we can technically do. Note that our FPP can still be quite low, even with such a low $k$ (Figure 4).

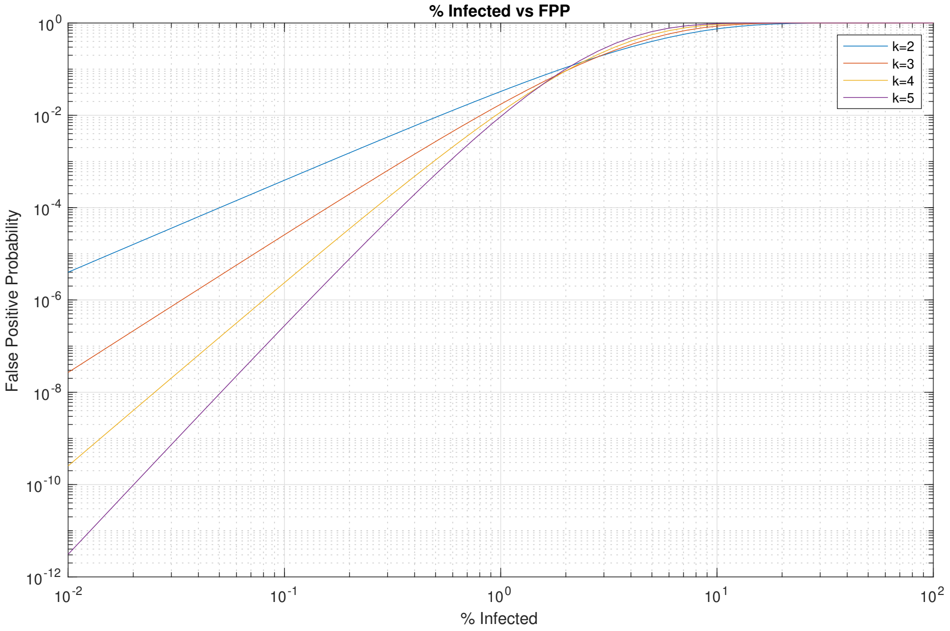


Figure 4: $FPP$ as % Infected varies with $m = 10^4$ and $b = 10^5$ and $k = 2, 3, 4$ or 5.

As an alternative approach, given an error threshold $\epsilon$ on the FPP, we can solve for a maximum compression factor $c = \frac{b}{m}$. This constraint is

$$c = \frac{-\ln(1 - \epsilon^{\frac{1}{k}})}{kp}.$$

This $c$ can be interpreted as the maximum number of samples per barcode we can run given an error threshold $\epsilon$, $k$ barcodes per sample, and some proportion $p$ infected. Let's choose $\epsilon = 0.0001$ or 0.01%. Then, we can examine the maximum theoretical compression factor for various $k$ as the % infected varies (Figure 5).

We can also calculate the max % infected that ensures that the error rate is still under threshold (Table 3). Note that scaling down the number of samples per batch by a factor $s$ also scales up the maximum percentage infected by a factor $s$, as the maximum % infected is inversely proportional to $b$ for a chosen error rate. For this regime, increasing $k$ will continue to raise the maximum % infected, before coming crashing down.

For $k = 4$, we can maintain a false positive probability of $< 0.01\%$ with $b = 10^5$ and $m = 10^4$ if the total % infected remains under 0.263%. If this is exceeded, we can scale down the number of samples per batch. If we reduce down to $b = 5 \times 10^4$, % infected needs to remain below 0.526% to maintain this error rate. Similarly, 1.317% max for $b = 2 \times 10^4$ and 2.634% max for $b = 10^4$. For $k = 5$, we can maintain a FPP $< 0.01\%$ with $b = 10^5$ and $m = 10^4$ if the total % infected remains under 0.345%.

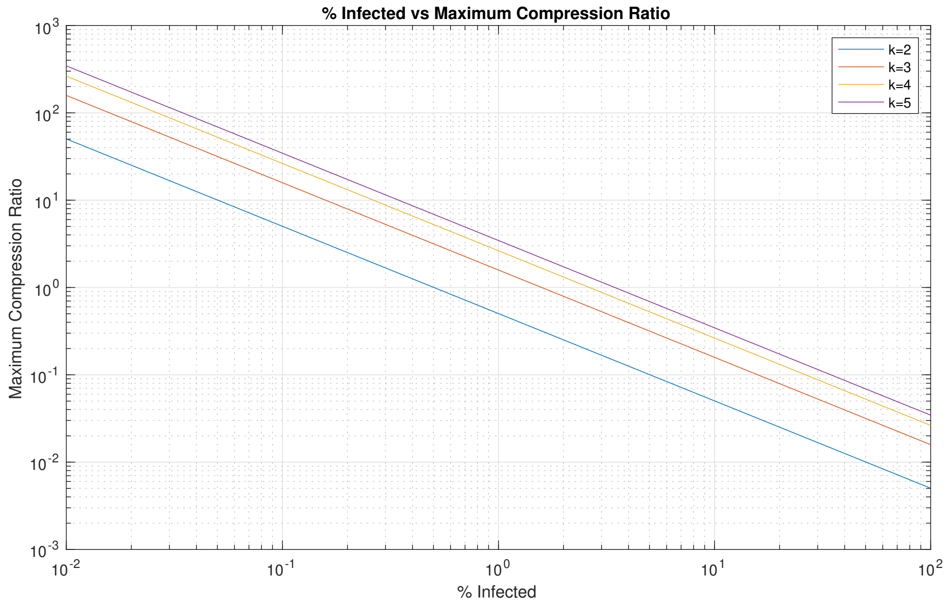


Figure 5: Maximum compression ratio as the % infected varies for $m = 10^4$ and $b = 10^5$.

| FPP (in %) | $k$ | % Infected Max ($b = 10^4$) | % Infected Max ($b = 10^5$) |
|---|---|---|---|
| 0.01 | 2 | 0.503 | 0.050 |
| 0.02 | 2 | 0.712 | 0.071 |
| 0.05 | 2 | 1.131 | 0.113 |
| 0.1 | 2 | 1.607 | 0.161 |
| 0.01 | 3 | 1.584 | 0.158 |
| 0.02 | 3 | 2.009 | 0.201 |
| 0.05 | 3 | 2.757 | 0.276 |
| 0.1 | 3 | 3.512 | 0.351 |
| 0.01 | 4 | 2.634 | 0.263 |
| 0.02 | 4 | 3.165 | 0.317 |
| 0.05 | 4 | 4.049 | 0.405 |
| 0.1 | 4 | 4.895 | 0.490 |
| 0.01 | 5 | 3.451 | 0.345 |
| 0.02 | 5 | 4.019 | 0.402 |
| 0.05 | 5 | 4.935 | 0.494 |
| 0.1 | 5 | 5.785 | 0.579 |

Table 3: Max % infected for various $k$ and FPPs

#### A.4 Single Bloom Filter with Barcode Loss

##### A.4.1 Modeling barcode loss

We will now model what happens if our query for whether a barcode is positive fails sometimes, considering the two types of errors from section 1. We will continue to assume that no negative barcodes will show up as positive.

The first type of error, where a given barcode never functions properly, occurs with probability $\Delta_{synth}$. In the Bloom filter model, for any specific bit in the bit array, this corresponds to having the bit remain at 0, regardless of the number of attempts at flipping it, with a probability of $\Delta_{synth}$.

The second type of error, where a given positive barcode may not be picked up for a particular sample, occurs with probability $\Delta_{stoch}$. In the Bloom filter model, for any specific bit in the bit array that works, this corresponds to having the bit not flip for a given attempt at flipping it with a probability of $\Delta_{stoch}$.

##### A.4.2 Error rates with barcode loss

We can also change the criteria with which we call a sample positive. In this first model, we will continue to only call a sample positive if all $k$ barcodes for the sample turn up positive. If at least one of the $k$ barcodes is negative, we will say it is negative.

We can calculate the False Positive Probability for this model ($\text{FPP}_\Delta$) as

$$FPP_\Delta = (1-\Delta_{synth})^k \left(1-\left(1-\frac{1-\Delta_{stoch}}{m}\right)^{kn}\right)^k \approx (1-\Delta_{synth})^k \left(1-e^{-(1-\Delta_{stoch})\frac{kn}{m}}\right)^k .$$

Note that non-zero $\Delta_{stoch}$ and $\Delta_{synth}$ only serve to lower this FPP compared to the original FPP with no errors, so the original FPP is an upper bound on the FPP incorporating errors. Since we don't know the exact values of $\Delta_{stoch}$ or $\Delta_{synth}$, it may be wise to proceed using the FPP derived with no errors. This would help ensure that models are robust regardless of the true values of the error rates.

Barcode errors also introduce false negatives, as there is some probability that at least one of the $k$ corresponding barcodes for a given sample will not show up as positive even if it is positive. The False Negative Probability ($\text{FNP}_\Delta$) for this model is

$$\begin{aligned} FNP_\Delta &= 1-(1-\Delta_{synth})^k \left(1-\Delta_{stoch}\left(1-\frac{1-\Delta_{stoch}}{m}\right)^{k(n-1)}\right)^k \\ &\approx 1-(1-\Delta_{synth})^k \left(1-\Delta_{stoch}e^{-(1-\Delta_{stoch})\frac{k(n-1)}{m}}\right)^k . \end{aligned}$$

Since this is dependent on $n$, this probability will vary with the % of the population infected. The $FNP_\Delta$ curves for $k = 2, 3, 4$, and 5 with various parameters are shown below (Figure 6).

For low % infected, a large component of the $FNP_\Delta$ is contributed by $\Delta_{stoch}$. As the % infected increases, this contribution drops, but may still dominate over $\Delta_{synth}$. An upper-bound on this probability, $FNP_{\Delta,max}$, is given by

$$FNP_{\Delta,max} = 1-(1-\Delta_{synth})^k (1-\Delta_{stoch})^k .$$

For $\Delta_{synth} = 0.01$ and $\Delta_{stoch} = 0.05$, this is 0.12 for $k = 2$, 0.17 for $k = 3$, 0.22 for $k = 4$, and 0.26 for $k = 5$. If our values for $\Delta_{synth}$ and $\Delta_{stoch}$ are ten-fold overestimates, then $FNP_{\Delta,max}$ would become 0.012 for $k = 2$, 0.018 for $k = 3$, 0.024 for $k = 4$, and 0.029 for $k = 5$.

In this regime of $k$, as $k$ increases, $FPP_\Delta$ will decrease and $FNP_\Delta$ will increase. So, there is a trade-off between false negatives and false positives as we vary $k$.

At the current values of % infected, $FNP_\Delta$ is much higher than $FPP_\Delta$. For a good diagnostic, we want to prioritize a low false negative probability, while still maintaining a low false positive probability. So, we will next try to model a few different strategies for reducing the false negative probability. The first two strategies will still be on a single batch tested once, and the remainder will involve a second barcode.

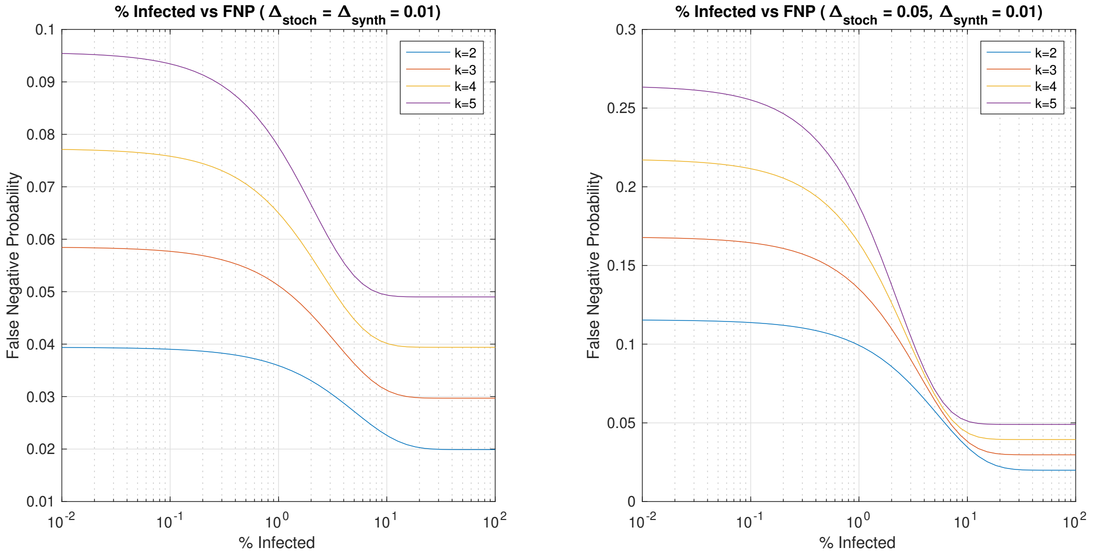


Figure 6: False Negative Probability ($FNP_{\Delta}$) as the % infected varies for $m = 10^4$ and $b = 10^5$.

##### A.4.3 Calling elements positive if $\geq k-1$ out of $k$ bits are positive

One potential strategy for augmenting a single Bloom filter with barcode errors is to change our criterion when we query whether a given element is in the set. We can do this by saying that an element is in the set if $\geq k-1$ out of the $k$ corresponding bits are 1. This is equivalent to calling a sample positive if $\geq k-1$ out of its $k$ corresponding barcodes are positive.

Under this scheme, we can calculate the error rates ($FNP_{\Delta 2}$ and $FPP_{\Delta 2}$) as

$$FPP_{\Delta 2} = FPP_{\Delta} + k\left(\Delta_{synth} + (1-\Delta_{synth})\left(1 - \frac{1-\Delta_{stoch}}{m}\right)^{kn}\right)$$
$$\left((1-\Delta_{synth})\left(1-\left(1-\frac{1-\Delta_{stoch}}{m}\right)^{kn}\right)\right)^{k-1}$$
$$FNP_{\Delta 2} = FNP_{\Delta} - k\left(\Delta_{synth} + (1-\Delta_{synth})\,\Delta_{stoch}\left(1-\frac{1-\Delta_{stoch}}{m}\right)^{k(n-1)}\right)$$
$$\left((1-\Delta_{synth})\left(1-\Delta_{stoch}\left(1-\frac{1-\Delta_{stoch}}{m}\right)^{k(n-1)}\right)\right)^{k-1},$$

derived in Appendix C along with exponential approximations. An upper bound on the false negative probability ($FNP_{\Delta 2,max}$) can be calculated as

$$FNP_{\Delta 2,max} = FNP_{\Delta,max} - k\left(\Delta_{synth} + (1-\Delta_{synth})\,\Delta_{stoch}\right)\left((1-\Delta_{synth})\,(1-\Delta_{stoch})\right)^{k-1}.$$

This upper bound on the false negative probability, for $\Delta_{synth} = 0.01$ and $\Delta_{stoch} = 0.05$, is 0.0035 for $k=2$, 0.010 for $k=3$, 0.020 for $k=4$, and 0.031 for $k=5$, much lower than before. A comparison between $FNP_{\Delta 2}$ and $FPP_{\Delta 2}$ and the original $FNP_{\Delta}$ and $FPP_{\Delta}$ is shown in Figure 7.

This strategy allows for a much lower false negative probability, at the cost of a higher false positive probability. With $k$ values that are so low, this might not be a reasonable approach, but it sets the stage for more complex strategies that can use this as part of their model.

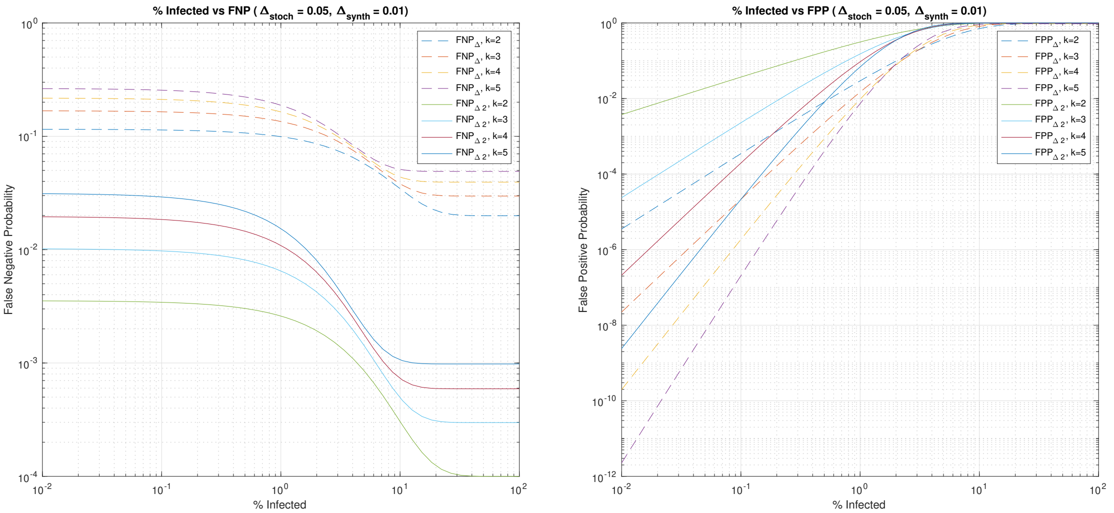


Figure 7: $FNP_\Delta$, $FPP_\Delta$, $FNP_{\Delta 2}$, $FPP_{\Delta 2}$ as the % infected varies for $m = 10^4$ and $b = 10^5$.

###### A.4.4 Calling elements positive if $\geq k'$ out of $k$ bits are positive

We can generalize to a strategy where we call a sample positive if $\geq k'$ out of the $k$ corresponding bits are 1. The error rates for this scheme, with $k' < k$, are

$$FPP_{\Delta k'} = FPP_\Delta + \sum_{i=k'}^{k-1} \binom{k}{i} \left(\Delta_{synth} + (1-\Delta_{synth})\left(1 - \frac{1-\Delta_{stoch}}{m}\right)^{kn}\right)^{i} \left((1-\Delta_{synth})\left(1-\left(1-\frac{1-\Delta_{stoch}}{m}\right)^{kn}\right)\right)^{k-i}$$

$$FNP_{\Delta k'} = FNP_\Delta - \sum_{i=k'}^{k} \binom{k}{i} \left(\Delta_{synth} + (1-\Delta_{synth})\,\Delta_{stoch}\left(1 - \frac{1-\Delta_{stoch}}{m}\right)^{k(n-1)}\right)^{k-i} \left((1-\Delta_{synth})\left(1-\Delta_{stoch}\left(1-\frac{1-\Delta_{stoch}}{m}\right)^{k(n-1)}\right)\right)^{i}.$$

derived in Appendix C along with exponential approximations and an upper bound $FNP_{\Delta k', max}$.

Continuing to lower $k'$ leads to further drops in the false negative probability, with further gains in the false positive probability. Since the impact of $k' = k-1$ on the false positive probability was already possibly too much, this strategy is not very reasonable if we only have a single Bloom filter with $b = 100,000$ and $m = 10,000$ with low $k$. This becomes more useful given a second set of barcodes.

##### A.5 Adding a Second Barcode with No Redundancy

There are many possible extensions if we have a set of $m_2$ barcode 2s available. Reasonable numbers for $m_2$ practically are 10, 96, or 384.

###### A.5.1 Pooling into non-overlapping sub-batches, no first barcodes

Suppose that we did not initially barcode the samples with the scheme described above, and were only given the $m_2$ orthogonal barcodes/wells. Then, assuming we can choose which samples go into which wells, this problem reduces to the initial problem of a single Bloom filter with $m = m_2$.

Since $m_2$ is low, the false positive probability is significantly elevated and the false negative probability is lower as the % infected varies. So, a large batch size is not reasonable in this scenario. The $\Delta_{synth}$ term would correspond to some wells consistently failing even when containing a positive sample, and the $\Delta_{stoch}$ term would correspond to a sample loading error or dropping out during downstream sequencing. So the results here are generalizable to other scenarios.

##### A.5.2 Pooling into non-overlapping sub-batches, with first barcodes

One potential strategy is to split the batch of $b$ samples into $m_2$ non-overlapping sub-batches, using a single barcode of the $m_2$ orthogonal barcodes for each sub-batch. Then, each sub-batch can be modeled as an individual Bloom filter. So, we now have $m_2$ separate non-overlapping Bloom filters, each with a batch size of $b_2 = \frac{b}{m_2}$.

As the other parameters do not change, we can compute the exact error probabilities for our three models for a single bloom filter by plugging in $n_2 = pb_2$ for $n$ into the formulas for $FPP$, $FNP_\Delta$, $FPP_\Delta$, $FNP_{\Delta 2}$, $FPP_{\Delta 2}$, $FNP_{\Delta k'}$, and $FPP_{\Delta k'}$.

Since our batch size has dropped by a factor of $m_2$, $FPP_\Delta$ will decrease. However, the $FNP_{\Delta,max}$ is not affected by varying the batch size, so this is not helpful towards our goal if we continue to call a sample positive only if all $k$ bits are positive.

Alternatively, if we use the $\geq k-1$ out of $k$ criterion, we can reduce the false positive probability while retaining a perhaps reasonable false negative probability. The error rates for the three different $m_2$ values are shown on the next page (Figure 8), using the $\geq k-1$ and exactly $k$ models. By utilizing these sub-batches, the $FPP_{\Delta 2}$ is lowered to something much more reasonable, so we are able to make use of the lower $FNP_{\Delta 2,max}$.

Perhaps the most reasonable approach is to use the $\geq k'$ out of $k$ criterion. Then, plugging in $n_2 = \frac{n}{m_2}$, the error rates are

$$FNP_{\Delta k',m_2} = 1 - \sum_{i=k'}^{k} \binom{k}{i} \left( \Delta_{synth} + (1-\Delta_{synth})\,\Delta_{stoch} \left(1 - \frac{1-\Delta_{stoch}}{m}\right)^{k\left(\frac{n}{m_2}-1\right)} \right)^{k-i}$$
$$\left( (1-\Delta_{synth}) \left( 1 - \Delta_{stoch} \left(1 - \frac{1-\Delta_{stoch}}{m}\right)^{k\left(\frac{n}{m_2}-1\right)} \right)\right)^{i}.$$

$$FPP_{\Delta k',m_2} = \sum_{i=k'}^{k} \binom{k}{i} \left( \Delta_{synth} + (1-\Delta_{synth}) \left(1 - \frac{1-\Delta_{stoch}}{m}\right)^{\frac{kn}{m_2}} \right)^{k-i}$$
$$\left( (1-\Delta_{synth}) \left( 1 - \left(1 - \frac{1-\Delta_{stoch}}{m}\right)^{\frac{kn}{m_2}} \right)\right)^{i}$$

Similarly, using our equations for the single Bloom filter with errors and the maximal FPP with no errors,

$$FNP_{\Delta k',m_2,max} = 1 - \sum_{i=k'}^{k} \binom{k}{i} \left(\Delta_{synth} + (1-\Delta_{synth})\,\Delta_{stoch}\right)^{k-i} \left((1-\Delta_{synth})(1-\Delta_{stoch})\right)^{i}.$$

$$FPP_{\Delta k',m_2,max} = \sum_{i=k'}^{k} \binom{k}{i} \left( \left(1 - \frac{1}{m}\right)^{\frac{kn}{m_2}} \right)^{k-i} \left( 1 - \left(1 - \frac{1}{m}\right)^{\frac{kn}{m_2}} \right)^{i}.$$

A scenario using this model is described in section 3.

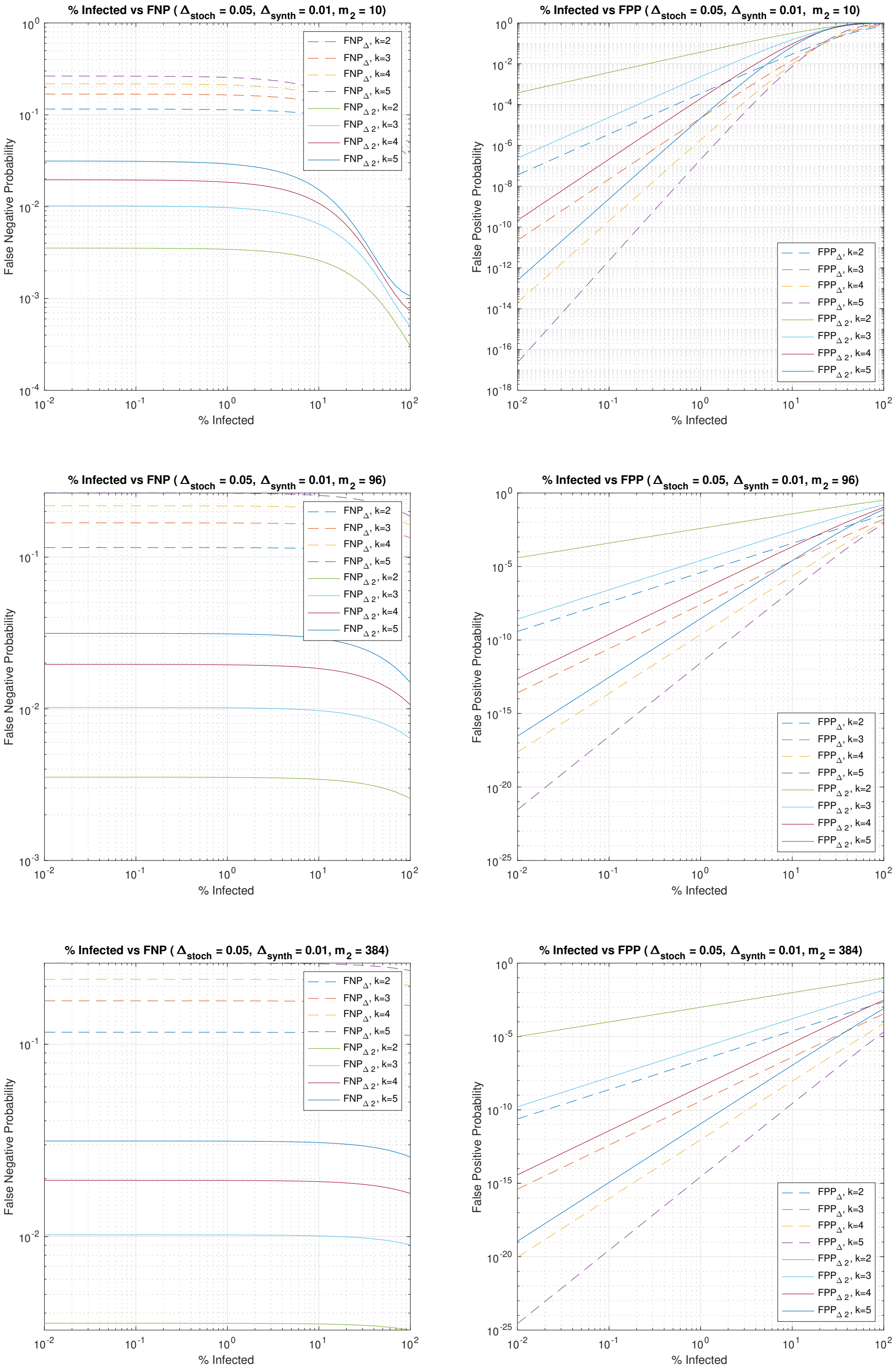


Figure 8: Error rates for various $m_2$ as the % infected varies for $m = 10^4$ and $b = 10^5$.

### B More Complex Scenarios

If a patient sample can be distributed into more than one sub-batch, then we can introduce redundancy across sub-batches. There are many potential theoretical scenarios where we can leverage this redundancy to improve our inference process. However, these scenarios may be challenging to perform in practice. Two of these scenarios, that have some potential for physical implementation, are described below.

#### B.1 Pooling into $m_2$ half-overlapping sub-batches

One other strategy would be to split the batch of $b$ samples into $m_2$ sub-batches, and then we dispense them into the $m_2$ orthogonally-barcoded wells in an overlapping fashion, so each well contains 2 different sub-batches. Then, each well will contain $\frac{2b}{m_2}$ samples, and each sample will be represented twice. An example of this is to split all of the samples into 10 batches, and then add batches 1 and 2 into well 1, batches 2 and 3 into well 2, and so forth.

This redundancy allows for a reduction of the false positive rate, and for various new criteria for calling a sample positive. Each well again functions as a separate Bloom filter, but the direct mathematical analysis is difficult here as the Bloom filters are no longer fully independent. We know that two wells containing a given sample have half of the samples that are the same. Since this independence assumption is violated, false negative and false positive probabilities may be best estimated for a given parameter set using numerical simulation.

#### B.2 Each sample goes into $k_2$ random wells

If we have a liquid handling robot, we could try to choose which wells a particular sample goes into. Taking a page from group testing, one way to use this to augment our testing is to have $k_2$ wells for each sample, where the design matrix is chosen such that each sample ends up in a unique combination of $k_2$ wells. For now, let $k_2 = 3$. This is possible since 384 choose 3 and 96 choose 3 are both big.

Since we have chosen unique combinations, the samples in any two wells are $\approx$ independent. This means that, for each sample, we now have $k_2$ different single Bloom filters with barcode errors containing it, each with $\frac{k_2 b}{m_2}$ samples. For $b = 100000$, $k_2 = 3$, $m_2 = 384$, this is $\approx 781$ samples per well.

Since the independence assumption is satisfied here, the error probabilities for a single one of these Bloom filters are the same as those computed in Appendix A.4.2, with a batch size of $\frac{k_2 b}{m_2}$. This approach opens the doors to new criteria for calling a sample positive, such as calling a sample positive if at least $k'$ out of $k$ barcodes are positive in 2 out of the 3 corresponding wells. The error rates for these kinds of approaches can be computed using combinatorics with the error rates for a single one of these Bloom filters.

### C Math Appendix

#### C.1 Single Bloom Filter, No Barcode Loss

##### C.1.1 False Positive Probability

We will now derive the false positive probability of the Bloom filter. This makes certain assumptions about the hash functions at play, namely that they generate a uniform random distribution across all inputs, and that the hash functions are independent. This is basically true for our samples as we can assign the $m$ unique barcodes across all samples randomly, although we would likely assign $k$ different unique barcodes per sample which is not accounted for here.

The probability that a certain bit $m'$ is not set to 1 during the addition of a single element due to a certain hash function is $1 - \frac{1}{m}$. If the hash functions don't have a significant correlation, then, after all $k$ hash functions have been used, this probability is

$$\Pr[m' = 0] = \left(1 - \frac{1}{m}\right)^k$$

for a single element addition. If we now add $n$ total elements, then

$$\Pr[m' = 0] = \left(1 - \frac{1}{m}\right)^{kn}$$

and

$$\Pr[m' = 1] = 1 - \left(1 - \frac{1}{m}\right)^{kn}.$$

Now, let's focus on an element $e$ that is not in the set. The $k$ hash functions map it to $k$ bits. So, the probability they are all 1, producing a false positive, is

$$FPP = \left(1 - \left(1 - \frac{1}{m}\right)^{kn}\right)^k. \tag{1}$$

##### C.1.2 Optimal choice of $k$

We can try to solve for an optimal choice of $k$ that minimizes the False Positive Probability for an ideal Bloom Filter. To simplify things, we will approximate $\left(1 - \frac{1}{m}\right)^{kn}$ as $e^{-\frac{kn}{m}}$. This is a valid approximation for large $m$, and quite reasonable for our $m$ of 10000. Then, for a fixed $m$ and $n$, we want

$$\min_k \left(1 - \left(1 - \frac{1}{m}\right)^{kn}\right)^k \approx \min_k \left(1 - e^{-\frac{kn}{m}}\right)^k.$$

Taking the natural log, this becomes

$$\min_k k \ln\left(1 - e^{-\frac{kn}{m}}\right).$$

First order conditions give

$$\ln\left(1 - e^{-\frac{kn}{m}}\right) + \frac{kn}{m}\frac{e^{-\frac{kn}{m}}}{1 - e^{-\frac{kn}{m}}} = 0.$$

Let $\frac{kn}{m} = \ln x$. Then,

$$\begin{aligned} 0 &= \ln\left(1 - e^{-\ln x}\right) + \ln x \frac{e^{-\ln x}}{1 - e^{-\ln x}} \\ &= \ln\left(1 - \frac{1}{x}\right) + \ln x \frac{\frac{1}{x}}{1 - \frac{1}{x}}, \end{aligned}$$

which solves to $x = 2$. So, the optimal choice of $k$ is

$$k_{OPT} = \ln(2)\frac{m}{n}. \tag{2}$$

At $k_{OPT}$, we have that

$$\begin{aligned} FPP_{OPT} &= \left(1 - e^{-k_{OPT}\frac{n}{m}}\right)^{k_{OPT}} \\ &= \left(1 - e^{-\ln(2)}\right)^{k_{OPT}} \\ &= 0.5^{k_{OPT}}. \end{aligned} \tag{3}$$

###### C.1.3 Maximal Compression Factor

We can also use our approximation to solve for a constraint on the $b$ to $m$ ratio given an error threshold $\epsilon$. Let $c$ be the compression factor $c = \frac{b}{m}$. Then,

$$\left(1 - e^{-\frac{kpb}{m}}\right)^k \leq \epsilon$$

or

$$e^{-kpc} \geq 1 - \epsilon^{\frac{1}{k}}$$

so

$$c \leq \frac{-\ln(1 - \epsilon^{\frac{1}{k}})}{kp}.$$

This $c$ we solve for can be interpreted as the number of samples per barcode we can run given the error threshold and some proportion $p$ infected. So, for a given choice of $p, k$, and $\epsilon$,

$$c_{max} = \frac{-\ln(1 - \epsilon^{\frac{1}{k}})}{kp}. \tag{4}$$

Rearranging, we also obtain that, for a given choice of $c, k$, and $\epsilon$,

$$p_{max} = \frac{-\ln(1 - \epsilon^{\frac{1}{k}})}{kc}. \tag{5}$$

#### C.2 Single Bloom Filter with Barcode Loss

##### C.2.1 $FPP_\Delta$ and $FNP_\Delta$

Given the two types of errors introduced above, we can attempt to calculate the false positive and false negative probabilities. We will continue to assume that the hash functions are independent, so we aren't accounting for the fact that the $k$ barcodes for a given sample are probably all different (except for one step in the $FNP_\Delta$ derivation).

Let's first calculate the false positive probability. For a given bit $m'$, that at least functions some portion of the time, the probability it is not set to 1 during the addition of a single element due to a certain hash function is $1 - (1 - \Delta_{stoch})\,\frac{1}{m}$. After all $k$ hash functions have been used, this probability is

$$\Pr[m' = 0] = \left(1 - \frac{1 - \Delta_{stoch}}{m}\right)^k$$

for a single element addition. If we now add $n$ total elements, then

$$\Pr[m' = 0] = \left(1 - \frac{1 - \Delta_{stoch}}{m}\right)^{kn}.$$

Incorporating the $\Delta_{synth}$ error,

$$\Pr[m' = 0] = \Delta_{synth} + (1 - \Delta_{synth}) \left(1 - \frac{1 - \Delta_{stoch}}{m}\right)^{kn}$$
$$\Pr[m' = 1] = (1 - \Delta_{synth}) \left(1 - \left(1 - \frac{1 - \Delta_{stoch}}{m}\right)^{kn}\right).$$

Now, let's focus on an element $e$ that is not in the set. The $k$ hash functions map it to $k$ bits. So, the probability they are all 1, producing a false positive, is

$$FPP_{\Delta} = (1 - \Delta_{synth})^k \left(1 - \left(1 - \frac{1 - \Delta_{stoch}}{m}\right)^{kn}\right)^k. \quad (6)$$

Note that we can write

$$1 - \left(1 - \frac{1 - \Delta_{stoch}}{m}\right)^{kn} = 1 - \left(\left(1 - \frac{1}{m/(1 - \Delta_{stoch})}\right)^{m/(1-\Delta_{stoch})}\right)^{\frac{1-\Delta_{stoch}}{m} kn}$$
$$\approx 1 - e^{-(1-\Delta_{stoch})\frac{kn}{m}}$$

for large $m$. This is a very close approximation for $m = 10000$. So, the false positive probability incorporating barcode errors, $FPP_{\Delta}$, is approximately given by

$$FPP_{\Delta} \approx (1 - \Delta_{synth})^k \left(1 - e^{-(1-\Delta_{stoch})\frac{kn}{m}}\right)^k. \quad (7)$$

We will now compute the false negative probability. Let's focus on an element $e'$ that is in the set. The $k$ hash functions map it to $k$ bits. The probability of a false negative is $1-$ (the probability it is called as positive), which requires all $k$ bits to be positive. For a single one of these $k$ bits $m$", it will be 0 with probability

$$\Pr[m" = 0] = \Delta_{synth} + (1 - \Delta_{synth}) \Delta_{stoch} \left(1 - \frac{1 - \Delta_{stoch}}{m}\right)^{k(n-1)},$$

assuming that the $k$ barcodes for $e'$ were different. Then, the probability that $m$" is 1 is

$$\Pr[m" = 1] = (1 - \Delta_{synth}) \left(1 - \Delta_{stoch} \left(1 - \frac{1 - \Delta_{stoch}}{m}\right)^{k(n-1)}\right).$$

Thus, the false negative probability incorporating barcode errors, $FNP_{\Delta}$, is

$$FNP_{\Delta} = 1 - (1 - \Delta_{synth})^k \left(1 - \Delta_{stoch} \left(1 - \frac{1 - \Delta_{stoch}}{m}\right)^{k(n-1)}\right)^k \quad (8)$$
$$\approx 1 - (1 - \Delta_{synth})^k \left(1 - \Delta_{stoch} e^{-(1-\Delta_{stoch})\frac{k(n-1)}{m}}\right)^k \quad (9)$$

by the same approximation as above.

Note that there is a dependence on the number of people infected $n$ in a given batch, due to the possibility of some other positive samples compensating for the dropped out barcode(s) for a given positive sample. We can produce an upper-bound on this probability, $FNP_{\Delta,max}$ by not allowing for this compensation, giving

$$FNP_{\Delta,max} = 1 - (1 - \Delta_{synth})^k (1 - \Delta_{stoch})^k. \quad (10)$$

##### C.2.2 Positives given as $\geq k-1$ out of $k$ barcodes positive

We've now established that for a random given bit $m'$, after $n$ elements have been inserted,

$$\Pr[m'=0] = \Delta_{synth} + (1-\Delta_{synth})\left(1-\frac{1-\Delta_{stoch}}{m}\right)^{kn}$$

$$\Pr[m'=1] = (1-\Delta_{synth})\left(1-\left(1-\frac{1-\Delta_{stoch}}{m}\right)^{kn}\right).$$

Now, let's focus on an element $e$ that is not in the set. The $k$ hash functions map it to $k$ bits. Under this model, at least $k-1$ bits have to be 1, so, the probability of a false positive, is

$$\begin{aligned} FPP_{\Delta 2} &= \Pr[m'=1]^k + k\Pr[m'=0]\Pr[m'=1]^{k-1} \\ &= FPP_{\Delta} + k\Pr[m'=0]\Pr[m'=1]^{k-1}. \end{aligned}$$

So,

$$\begin{aligned} FPP_{\Delta 2} = FPP_{\Delta} &+ k\left(\Delta_{synth} + (1-\Delta_{synth})\left(1-\frac{1-\Delta_{stoch}}{m}\right)^{kn}\right) \\ &\left((1-\Delta_{synth})\left(1-\left(1-\frac{1-\Delta_{stoch}}{m}\right)^{kn}\right)\right)^{k-1} \\ = &\left((1-\Delta_{synth})\left(1-\left(1-\frac{1-\Delta_{stoch}}{m}\right)^{kn}\right)\right)^{k} \\ &+ k\left(\Delta_{synth} + (1-\Delta_{synth})\left(1-\frac{1-\Delta_{stoch}}{m}\right)^{kn}\right) \\ &\left((1-\Delta_{synth})\left(1-\left(1-\frac{1-\Delta_{stoch}}{m}\right)^{kn}\right)\right)^{k-1}. \end{aligned} \tag{11}$$

Using our approximation,

$$\begin{aligned} FPP_{\Delta 2} \approx FPP_{\Delta} &+ k\left(\Delta_{synth} + (1-\Delta_{synth})\, e^{-(1-\Delta_{stoch})\frac{k(n-1)}{m}}\right) \\ &\left((1-\Delta_{synth})\left(1-e^{-(1-\Delta_{stoch})\frac{k(n-1)}{m}}\right)\right)^{k-1} \\ \approx &\left((1-\Delta_{synth})\left(1-e^{-(1-\Delta_{stoch})\frac{kn}{m}}\right)\right)^{k} + k\left(\Delta_{synth} + (1-\Delta_{synth})\, e^{-(1-\Delta_{stoch})\frac{k(n-1)}{m}}\right) \\ &\left((1-\Delta_{synth})\left(1-e^{-(1-\Delta_{stoch})\frac{k(n-1)}{m}}\right)\right)^{k-1}. \end{aligned} \tag{12}$$

We will now compute the false negative probability. Let's focus on an element $e'$ that is in the set. The $k$ hash functions map it to $k$ bits. The probability of a false negative is 1−(the probability it is called as positive), which requires $\geq k-1$ bits to be positive. For a single one of these $k$ bits $m$", we have

$$\Pr[m"=0] = \Delta_{synth} + (1-\Delta_{synth})\,\Delta_{stoch}\left(1-\frac{1-\Delta_{stoch}}{m}\right)^{k(n-1)}$$

$$\Pr[m"=1] = (1-\Delta_{synth})\left(1-\Delta_{stoch}\left(1-\frac{1-\Delta_{stoch}}{m}\right)^{k(n-1)}\right).$$

So, the false negative probability incorporating barcode errors, $FNP_{\Delta 2}$, is

$$\begin{aligned} FNP_{\Delta 2} &= 1 - \Pr[m"=1]^k - k\Pr[m"=0]\Pr[m"=1]^{k-1} \\ &= FNP_{\Delta} - k\Pr[m"=0]\Pr[m"=1]^{k-1}. \end{aligned}$$

Expanding,

$$
\begin{aligned}
FNP_{\Delta 2} = FNP_{\Delta} - k &\left(\Delta_{synth} + (1-\Delta_{synth})\,\Delta_{stoch}\left(1-\frac{1-\Delta_{stoch}}{m}\right)^{k(n-1)}\right) \\
&\left((1-\Delta_{synth})\left(1-\Delta_{stoch}\left(1-\frac{1-\Delta_{stoch}}{m}\right)^{k(n-1)}\right)\right)^{k-1} \\
= 1 - (1-\Delta_{synth})^{k} &\left(1-\Delta_{stoch}\left(1-\frac{1-\Delta_{stoch}}{m}\right)^{k(n-1)}\right)^{k} \\
- k &\left(\Delta_{synth} + (1-\Delta_{synth})\,\Delta_{stoch}\left(1-\frac{1-\Delta_{stoch}}{m}\right)^{k(n-1)}\right) \\
&\left((1-\Delta_{synth})\left(1-\Delta_{stoch}\left(1-\frac{1-\Delta_{stoch}}{m}\right)^{k(n-1)}\right)\right)^{k-1}.
\end{aligned} \tag{13}
$$

Using our approximation,

$$
\begin{aligned}
FNP_{\Delta 2} \approx FNP_{\Delta} - k &\left(\Delta_{synth} + (1-\Delta_{synth})\,\Delta_{stoch} e^{-(1-\Delta_{stoch})\frac{k(n-1)}{m}}\right) \\
&\left((1-\Delta_{synth})\left(1-\Delta_{stoch} e^{-(1-\Delta_{stoch})\frac{k(n-1)}{m}}\right)\right)^{k-1} \\
\approx 1 - (1-\Delta_{synth})^{k} &\left(1-\Delta_{stoch} e^{-(1-\Delta_{stoch})\frac{k(n-1)}{m}}\right)^{k} \\
- k &\left(\Delta_{synth} + (1-\Delta_{synth})\,\Delta_{stoch} e^{-(1-\Delta_{stoch})\frac{k(n-1)}{m}}\right) \\
&\left((1-\Delta_{synth})\left(1-\Delta_{stoch} e^{-(1-\Delta_{stoch})\frac{k(n-1)}{m}}\right)\right)^{k-1}.
\end{aligned} \tag{14}
$$

Applying the same logic as above, we can compute an upper bound on this probability

$$
\begin{aligned}
FNP_{\Delta 2,max} &= FNP_{\Delta,max} - k\left(\Delta_{synth} + (1-\Delta_{synth})\,\Delta_{stoch}\right)\left((1-\Delta_{synth})(1-\Delta_{stoch})\right)^{k-1} \\
&= 1 - (1-\Delta_{synth})^{k}(1-\Delta_{stoch} e)^{k} \\
&\quad - k\left(\Delta_{synth} + (1-\Delta_{synth})\,\Delta_{stoch}\right)\left((1-\Delta_{synth})(1-\Delta_{stoch})\right)^{k-1}.
\end{aligned} \tag{15}
$$

##### C.2.3 Positives given as $\geq k'$ out of $k$ barcodes positive

First we will compute the false positive probability. Let's again focus on an element $e$ that is not in the set. The $k$ hash functions map it to $k$ bits. Under this model, at least $k'$ bits have to be 1, so, the probability of a false positive, is

$$
\begin{aligned}
FPP_{\Delta k'} &= \sum_{i=k'}^{k}\binom{k}{i}\Pr[m'=1]^{i}\Pr[m'=0]^{k-i} \\
&= FPP_{\Delta} + \sum_{i=k'}^{k-1}\binom{k}{i}\Pr[m'=1]^{i}\Pr[m'=0]^{k-i}
\end{aligned}
$$

for $k' < k$.

So,

$$
\begin{aligned}
FPP_{\Delta k'} = & \sum_{i=k'}^{k} \binom{k}{i} \left( \Delta_{synth} + (1-\Delta_{synth}) \left(1 - \frac{1-\Delta_{stoch}}{m}\right)^{kn} \right)^{k-i} \\
& \left( (1-\Delta_{synth}) \left(1 - \left(1 - \frac{1-\Delta_{stoch}}{m}\right)^{kn}\right)\right)^{i} \\
= & FPP_{\Delta} + \sum_{i=k'}^{k-1} \binom{k}{i} \left( \Delta_{synth} + (1-\Delta_{synth}) \left(1 - \frac{1-\Delta_{stoch}}{m}\right)^{kn} \right)^{k-i} \\
& \left( (1-\Delta_{synth}) \left(1 - \left(1 - \frac{1-\Delta_{stoch}}{m}\right)^{kn}\right)\right)^{i}.
\end{aligned} \tag{16}
$$

Using our approximation,

$$
\begin{aligned}
FPP_{\Delta k'} \approx & FPP_{\Delta} + \sum_{i=k'}^{k-1} \binom{k}{i} \left( \Delta_{synth} + (1-\Delta_{synth}) e^{-(1-\Delta_{stoch})\frac{k(n-1)}{m}} \right)^{k-i} \\
& \left( (1-\Delta_{synth}) \left(1 - e^{-(1-\Delta_{stoch})\frac{k(n-1)}{m}}\right)\right)^{i} \\
\approx & \sum_{i=k'}^{k} \binom{k}{i} \left( \Delta_{synth} + (1-\Delta_{synth}) e^{-(1-\Delta_{stoch})\frac{k(n-1)}{m}} \right)^{k-i} \\
& \left( (1-\Delta_{synth}) \left(1 - e^{-(1-\Delta_{stoch})\frac{k(n-1)}{m}}\right)\right)^{i}.
\end{aligned} \tag{17}
$$

We will now compute the false negative probability. Let's focus on an element $e'$ that is in the set. The $k$ hash functions map it to $k$ bits. The probability of a false negative is $1-$(the probability it is called as positive), which requires $\geq k'$ bits to be positive. So, the false negative probability incorporating barcode errors, $FNP_{\Delta 2}$, is

$$
\begin{aligned}
FNP_{\Delta k'} &= 1 - \sum_{i=k'}^{k} \binom{k}{i} \Pr[m\text{''} = 1]^{i} \Pr[m\text{''} = 0]^{k-1} \\
&= FNP_{\Delta} - \sum_{i=k'}^{k-1} \binom{k}{i} \Pr[m\text{''} = 1]^{i} \Pr[m\text{''} = 0]^{k-1}
\end{aligned}
$$

for $k' < k$. Expanding,

$$
\begin{aligned}
FNP_{\Delta k'} = & 1 - \sum_{i=k'}^{k} \binom{k}{i} \left( \Delta_{synth} + (1-\Delta_{synth}) \Delta_{stoch} \left(1 - \frac{1-\Delta_{stoch}}{m}\right)^{k(n-1)} \right)^{k-i} \\
& \left( (1-\Delta_{synth}) \left(1 - \Delta_{stoch}\left(1 - \frac{1-\Delta_{stoch}}{m}\right)^{k(n-1)}\right)\right)^{i} \\
= & FNP_{\Delta} - \sum_{i=k'}^{k} \binom{k}{i} \left( \Delta_{synth} + (1-\Delta_{synth}) \Delta_{stoch} \left(1 - \frac{1-\Delta_{stoch}}{m}\right)^{k(n-1)} \right)^{k-i} \\
& \left( (1-\Delta_{synth}) \left(1 - \Delta_{stoch}\left(1 - \frac{1-\Delta_{stoch}}{m}\right)^{k(n-1)}\right)\right)^{i}.
\end{aligned} \tag{18}
$$

Using our approximation,

$$\begin{aligned} FNP_{\Delta k'} &\approx FNP_{\Delta} - \sum_{i=k'}^{k-1} \binom{k}{i} \left( \Delta_{synth} + \left(1 - \Delta_{synth}\right) \Delta_{stoch} e^{-\left(1-\Delta_{stoch}\right) \frac{k(n-1)}{m}} \right)^{k-i} \\ &\quad \left( \left(1 - \Delta_{synth}\right) \left(1 - \Delta_{stoch} e^{-\left(1-\Delta_{stoch}\right) \frac{k(n-1)}{m}} \right) \right)^{i} \\ &\approx 1 - \sum_{i=k'}^{k} \binom{k}{i} \left( \Delta_{synth} + \left(1 - \Delta_{synth}\right) \Delta_{stoch} e^{-\left(1-\Delta_{stoch}\right) \frac{k(n-1)}{m}} \right)^{k-i} \\ &\quad \left( \left(1 - \Delta_{synth}\right) \left(1 - \Delta_{stoch} e^{-\left(1-\Delta_{stoch}\right) \frac{k(n-1)}{m}} \right) \right)^{i}. \end{aligned} \tag{19}$$

Applying the same logic as above, we can compute an upper bound on this probability

$$\begin{aligned} FNP_{\Delta k', max} &= FNP_{\Delta, max} - \sum_{i=k'}^{k-1} \binom{k}{i} \left( \Delta_{synth} + \left(1 - \Delta_{synth}\right) \Delta_{stoch} \right)^{k-i} \\ &\quad \left( \left(1 - \Delta_{synth}\right) \left(1 - \Delta_{stoch}\right) \right)^{i} \\ &= 1 - \sum_{i=k'}^{k} \binom{k}{i} \left( \Delta_{synth} + \left(1 - \Delta_{synth}\right) \Delta_{stoch} \right)^{k-i} \\ &\quad \left( \left(1 - \Delta_{synth}\right) \left(1 - \Delta_{stoch}\right) \right)^{i}. \end{aligned} \tag{20}$$
