## Supplementary Note 2 for "LAMP-Seq: Population-Scale COVID-19 Diagnostics Using Combinatorial Barcoding"

#### LAMP-Seq: Population-scale COVID-19 Diagnostics Using Combinatorial Barcoding - Dual Barcoding

Building on Supplementary Note 1, we now consider an alternative form of barcoding, where patient samples can be barcoded with a combination of barcoded FIP and barcoded BIP primers. We model various scenarios with this dual barcoding scheme, characterize error rates, introduce a template switching error term $\Delta_{switch}$, and simulate realistic parameter sets.

Our overall goal here is to design a setup that achieves false negative probabilities (FNPs) and false positive probabilities (FPPs) of $< 0.2\%$ using a minimal number of total barcodes given parameters of 10 sub-batches per run, 1% positive patient samples, reasonable error rates ($\Delta_{stoch} = 0.05, \Delta_{synth} = 0, \Delta_{switch} = 0.02$), and either 100,000 or 1,000,000 patients per batch.

There are three barcodes available: patient samples will be individually barcoded at the RT-LAMP stage with a first set of barcodes (barcode 1 – FIP, barcode 2 - BIP), and then groups of patient samples will be barcoded with an additional set of orthogonal barcodes (barcode 3) in the process of preparing samples for Illumina sequencing. After sequencing, a given combination of the three barcodes can be called as positive or negative for SARS-CoV-2 viral RNA.

Let there be $m_1$ total unique barcode 1s, $m_2$ total unique barcode 2s, and $m_3$ total unique barcode 3s. Each patient sample receives $k_1$ pre-assigned barcode 1s and $k_2$ pre-assigned barcode 2s. All other notation remains the same as Supplementary Note 1. Each barcode 1 or barcode 2 is assumed to fail globally with probability $\Delta_{synth}$, and a specific barcode 1 - barcode 2 pair for a patient sample is assumed to fail with probability $\Delta_{stoch}$. There are three scenarios considered below: one where $k_1 = k_2 = 1$, one where $k_1 > 1$ and $k_2 = 1$, and one where $k_1, k_2 > 1$.

#### 1 Scenario 3: $k_1 = k_2 = 1$

If $b < m_1 \cdot m_2 \cdot m_3$, then every sample in a batch can be assigned a unique barcode. For testing $b = 100,000$ samples, 100 barcode 1s, 100 barcode 2s, and 10 barcode 3s would suffice. Testing $b = 1,000,000$ samples would only require an increase in the number of barcode 3s to 100. Then, every patient sample in a batch would have a unique barcode 1 – barcode 2 - barcode 3 group.

It is unlikely, although not impossible, for $N < m_1 \cdot m_2 \cdot m_3$, which would enable a unique barcode for every sample from the total population. This suggests that batches would have to be defined in some way, and that this scenario is better suited for synchronous testing.

Since each patient has a unique barcode group, there are no false positives. However, with barcode loss, there may be false negatives. Each patient sample now has two barcodes in the RT-LAMP reaction, so the overall false negative probability is $1 - (1 - \Delta_{stoch})(1 - \Delta_{synth})^2$.

#### 2 Scenario 4: $k_1 > 1$, $k_2 = 1$

With a liquid handler, it may be possible to give each patient sample $k_1$ different barcode 1s, with $k_1 > 1$, and a single barcode 2 in a pre-assigned way. Then, if $N < \binom{m_1}{k_1} \cdot m_2$, every patient sample for the entire population could have a unique combination of barcode 1s and barcode 2s. This does not mean that every barcode 1 – barcode 2 pair would correspond to a unique patient sample, as this is only possible if $N < \frac{m_1 m_2}{k_1}$.

For this scenario, we imagine an asynchronous sample collection system where $b$ of these samples, as they come in, are split into $m_3$ sub-batches. Each sub-batch then gets an additional unique barcode 3. Patient samples are inferred as positive if, after sequencing, $\geq k'_{12}$ out of the $k_1$ corresponding barcode 1 – barcode 2 - barcode 3 groups are positive.

This is similar to scenario 2 in Supplementary Note 1, except that the barcode 2s here provides a psuedo-sub-batch barcode without requiring physical distribution of samples. We assume that the number of barcode 2s, $m_2$, is small enough that it is possible to validate each barcode, so we do not consider $\Delta_{synth}$ errors for barcode 2s. Combining this with the barcode 3s, we effectively have $m_2 \cdot m_3$ separate non-overlapping sub-pools with $\frac{b}{m_2 m_3}$ samples each.

Since the barcode 1 – barcode 2 pairs no longer correspond to unique patient samples, false positives are possible using this approach. False negatives are also still possible, as a positive sample could be inferred as a negative sample if more than $k_1 - k'_{12}$ of the $k_1$ corresponding barcode groups are lost. Using a modified Bloom filter model (see Appendix A, Supplementary Note 1, and (1)), we can compute the false positive probability $\left(FPP_{\Delta k'_{12},4}\right)$ and false negative probability $\left(FNP_{\Delta k'_{12},4}\right)$ for this approach as

$$FPP_{\Delta k'_{12},4} = \sum_{i=k'_{12}}^{k_1} \binom{k_1}{i} \left(\Delta_{synth} + (1-\Delta_{synth})\left(1 - \frac{1-\Delta_{stoch}}{m_1}\right)^{\frac{k_1 n}{m_2 m_3}}\right)^{k_1 - i}$$
$$\left((1-\Delta_{synth})\left(1 - \left(1 - \frac{1-\Delta_{stoch}}{m_1}\right)^{\frac{k_1 n}{m_2 m_3}}\right)\right)^{i}$$

$$FNP_{\Delta k'_{12},4} = 1 - \sum_{i=k'_{12}}^{k_1} \binom{k_1}{i} \left(\Delta_{synth} + (1-\Delta_{synth})\,\Delta_{stoch}\left(1 - \frac{1-\Delta_{stoch}}{m_1}\right)^{k_1\left(\frac{n}{m_2 m_3}-1\right)}\right)^{k_1 - i}$$
$$\left((1-\Delta_{synth})\left(1 - \Delta_{stoch}\left(1 - \frac{1-\Delta_{stoch}}{m_1}\right)^{k_1\left(\frac{n}{m_2 m_3}-1\right)}\right)\right)^{i}.$$

These expressions are similar to those in Scenario 2, and analyses there apply here as well.

### 3 Scenario 5: $k_1, k_2 > 1$

It may also be possible to give each patient sample $k_1$ different barcode 1s and $k_2$ different barcode 2s, with $k_1, k_2 > 1$, in a pre-assigned way. Then, if $N < \binom{m_1}{k_1} \cdot \binom{m_2}{k_2}$, every patient sample for the entire population could have a unique combination of barcode 1s and barcode 2s. This does not mean that every barcode 1 – barcode 2 pair would correspond to a unique patient sample, as this is only possible if $N < \frac{m_1 m_2}{k_1 k_2}$.

The RT-LAMP reaction randomly incorporates FIP and BIP primers during amplification, so a positive patient sample can produce all $k_1 \cdot k_2$ barcode pairs. As an example, consider $k_1 = k_2 = 3$. We would then introduce 3 barcoded FIP primers and 3 barcoded BIP primers per RT-LAMP reaction, with a result of 9 distinct barcode 1 – barcode 2 pairs per sample.

We imagine a asynchronous sample collection system for this scenario, similar to scenario 4, and will consider 2 schemes for decoding which patients are positive. In the first decoder scheme, a patient sample is inferred to be positive, if, after sequencing, at least $k'_2$ of the $k_2$ patient-specific barcode 2s contain at least $k'_1$ of the $k_1$ patient-specific barcode 1 – barcode 2 pairs. In the second decoder scheme, a patient sample is inferred to be positive if, after sequencing, $\geq k'_{12}$ out of the $k_1 \cdot k_2$ patient-specific barcode 1 – barcode 2 - barcode 3 groups come up as positive.

This is similar to scenario 2 in Supplementary Note 1, with physical placement in a combination of $k_2$ wells corresponding to the combination of $k_2$ primers per sample here. The random barcode selection among samples allows us to calculate approximate error rates. We assume that the number of barcode 2s, $m_2$, is small enough that it is possible to validate each barcode, so we do not model $\Delta_{synth}$ errors for barcode 2s.

The FPP and FNP of the first decoder scheme can be approximated using equations from Supplementary Note 1; we exclude it here. Under the second decoder scheme, the false positive probability $\left(FPP_{\Delta k'_{12},5}\right)$ and false negative probability $\left(FNP_{\Delta k'_{12},5}\right)$ are approximately given by

$$FPP_{\Delta k'_{12},5} = \sum_{i=k'_{12}}^{k_1 k_2} \binom{k_1 k_2}{i} \left(\Delta_{synth} + (1-\Delta_{synth}) \left(1 - \frac{1-\Delta_{stoch}}{m_1}\right)^{\frac{k_1 k_2 n}{m_2 m_3}}\right)^{k_1 k_2 - i} \left((1-\Delta_{synth}) \left(1 - \left(1 - \frac{1-\Delta_{stoch}}{m_1}\right)^{\frac{k_1 k_2 n}{m_2 m_3}}\right)\right)^{i}$$

$$FNP_{\Delta k'_{12},5} = -\sum_{i=k'_{12}}^{k_1 k_2} \binom{k_1 k_2}{i} \left(\Delta_{synth} + (1-\Delta_{synth}) \Delta_{stoch} \left(1 - \frac{1-\Delta_{stoch}}{m_1}\right)^{k_1 \left(\frac{k_2 n}{m_2 m_3} - 1\right)}\right)^{k_1 k_2 - i} \left((1-\Delta_{synth}) \left(1 - \Delta_{stoch} \left(1 - \frac{1-\Delta_{stoch}}{m_1}\right)^{k_1 \left(\frac{k_2 n}{m_2 m_3} - 1\right)}\right)\right)^{i} + 1.$$

As the number of positive samples per batch $n$ increases, the false positive probability increases and the false negative probability decreases. We can compute an upper bound on the false negative probability $\left(FNP_{\Delta k'_{12},5,max}\right)$ that avoids this dependence as

$$FNP_{\Delta k'_{12},5,max} = -\sum_{i=k'_{12}}^{k_1 k_2} \binom{k_1 k_2}{i} \left(\Delta_{synth} + (1-\Delta_{synth}) \Delta_{stoch}\right)^{k_1 k_2 - i} \left((1-\Delta_{synth})(1-\Delta_{stoch})\right)^{i} + 1.$$

Overall, a lower $k'_{12}$ for a fixed $k_1 \cdot k_2$ leads to a reduced false negative probability at the cost of an increased false positive probability. The effect of varying $k_1$ and $k_2$ depends on the chosen $k'_{12}$ and on the proportion of the population infected $p$.

### 4 Modeling Template Switching

Under this dual barcoding scheme, template switching errors may occur during the PCR before sequencing. These are errors that lead to barcode swapping/index swapping between barcode 1 – barcode 2 pairs. Initial experiments have shown that the residual primers of negative samples do not lead to noticeable amounts of template swapping, likely due to sample dilution, so all template switching is assumed to occur between barcode 1 – barcode 2 pairs of positive samples.

As an example, consider two positive patient samples: one with barcode 1 – barcode 2 pair A–B and one with barcode 1 – barcode 2 pair C–D. Template switching would produce A–D and B–D as products. We model each template switching product as occurring with probability $\Delta_{switch}$. This probability has not been well characterized. We chose an initial value of 0.02 for simulations.

To provide some intuition, we switch to a graph-theoretic perspective. The barcode 1 – barcode 2 pairs can be modeled as edges on a bipartite graph formed by the set of barcode 1s ($U$, cardinality $m_1$) and the set of barcode 2s ($V$, cardinality $m_2$). Each patient sample is assigned a subset of cardinality $k_1$ from $U$ and a subset of cardinality $k_2$ from $V$. If a particular barcode 1 – barcode 2 pair is positive, the corresponding edge is part of the graph.

Template switching products then correspond to edges between any vertex in $U$ with degree $\geq 1$ and any vertex in $V$ with degree $\geq 1$. Each of these edges is added to the graph with probability $\Delta_{switch}$. Inference for patient samples is performed on the final graph once all template switching edges have been considered. The status of a particular patient sample is inferred by considering the edges on the induced sub-graph formed by the patient sample's specific subsets. Examples of this for scenario 3 and scenario 5 are shown in Figures 1 and 2.

Each template switching product in scenario 3 leads to a false negative, as each patient sample corresponds to a single barcode 1 – barcode 2 pair, which may be a concern. Scenarios 4 and 5 are able to mitigate this by requiring more than one positive barcode 1 – barcode 2 pair per sample. However, the number of possible template switching products and the number of template switching products formed also increases in scenario 4 and 5. So, when positive samples are sparse, scenarios 4 and 5 may perform better than scenario 3, but when positive samples are very common, barcode saturation occurs and scenario 3 outperforms scenario 4 and 5.

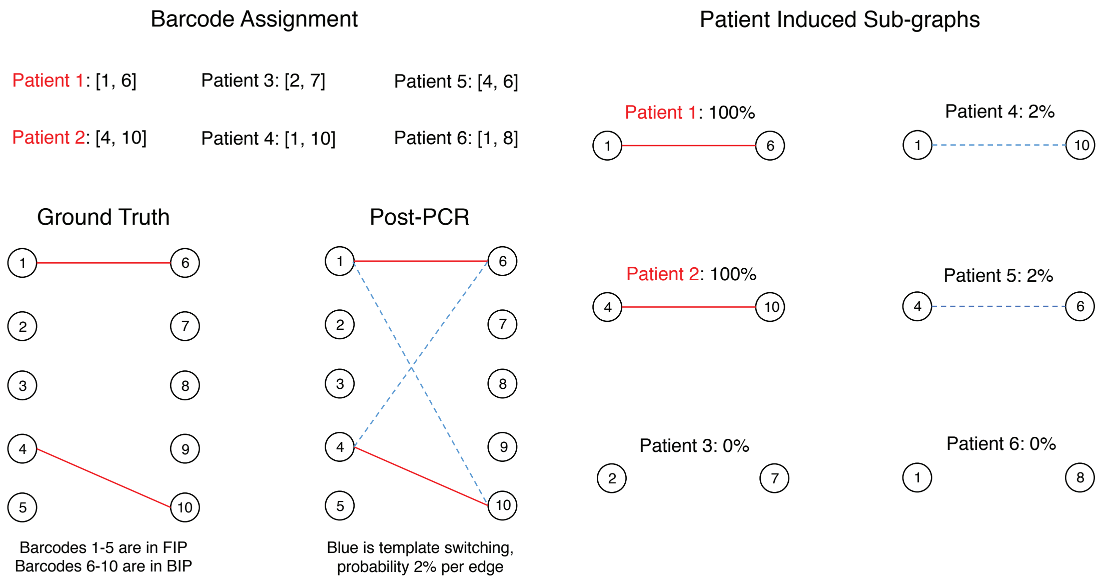


Figure 1: Template switching for scenario 3 with $m_1 = m_2 = 5$, $k_1 = k_2 = 1$, $\Delta_{switch} = 0.02$, and no barcode loss. The probability of inferring a particular sample as positive is shown next to each patient sample.

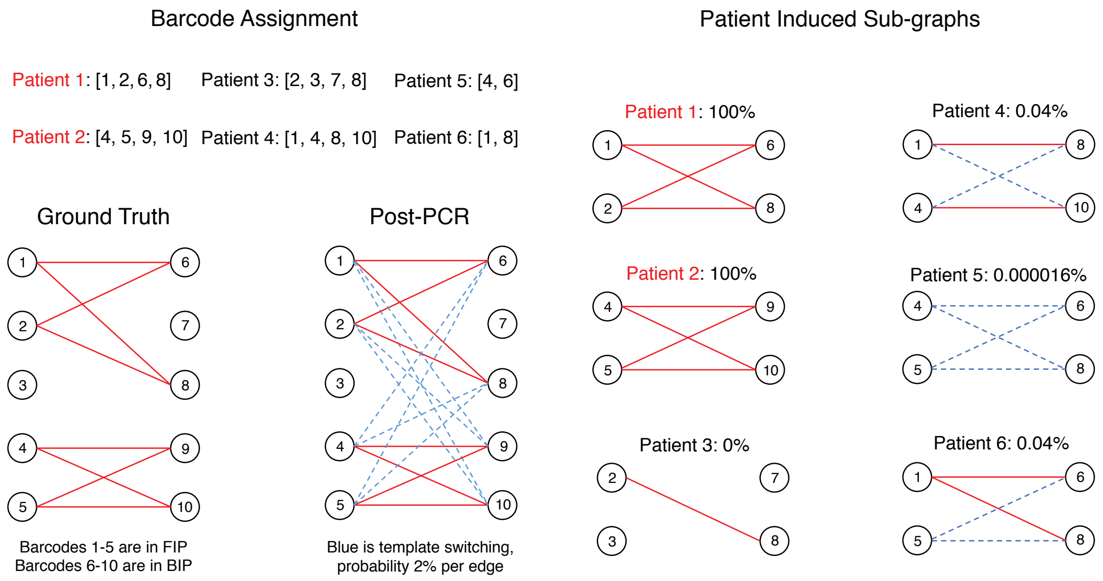


Figure 2: Template switching for scenario 5 with $m_1 = m_2 = 5$, $k_1 = k_2 = 2$, $\Delta_{switch} = 0.02$, $k'_{12} = 4$, and no barcode loss. The probability of inferring a particular sample as positive is shown next to each patient sample.

### 5 Numerical Simulations

Combining barcode loss and template switching, we ran numerical simulations to evaluate the performance of each scenario towards achieving our design goal. We chose $m_3 = 10$, so 10 sub-batches per run. Simulations were run for 5000 or 500 iterations, for $100,000$ samples per batch and $1,000,000$ samples per batch respectively.

We assumed that the number of barcodes used was small enough for physical validation of each barcode, so we set $\Delta_{synth} = 0$. The other error rates used were $\Delta_{stoch} = 0.05$ and $\Delta_{switch} = 0.02$, and we assumed that 1% of patient samples were positive.

#### 5.1 100,000 patient samples per batch

Our initial design goal was to achieve FNPs and FPPs of $< 0.2\%$ with a minimal number of total barcodes at 1% positive patient samples. With a small number of barcodes, scenario 3 ($m_1 = m_2 = 100$) and scenario 4 ($m_1 = m_2 = 96$, $k_1 = 5$, $k'_{12} = 3$) do not achieve sufficient error rates.

Scenario 5 with $m_1 = m_2 = 96$, $k_1 = k_2 = 3$, under the second decoder scheme with $k'_{12} = 6$ satisfies the design goal, suggesting that we can efficiently test $100,000$ patient samples with a total of $96 + 96 + 10 = 202$ barcodes. Increasing the number of barcode 1s and barcode 2s to 192 or 384 with $k'_{12} = 5$ lowers error rates further.

| Scenario | $m_1, m_2$ | $k_1$ | $k_2$ | $k'_{12}$ | $k'_1$ | $k'_2$ | Average FNP | Average FPP |
|---|---|---|---|---|---|---|---|---|
| 3 | 100 | 1 | 1 | – | – | – | 0.049864 | 0.0075493 |
| 4 | 96 | 5 | 1 | 3 | – | – | 0.000808 | 0.0059579 |
| 5 | 96 | 3 | 3 | – | 2 | 2 | 0.000168 | 0.0041614 |
| 5 | 96 | 3 | 3 | 6 | – | – | 0.000448 | 0.0001958 |
| 5 | 192 | 3 | 3 | 5 | – | – | 0.000104 | 0.0000206 |
| 5 | 384 | 3 | 3 | 5 | – | – | 0.000104 | 0.0000004 |

Table 1: FPP/FNP with $b = 100,000$, $m_3 = 10$, and $p = 0.01$ over 5000 iterations.

#### 5.2 1,000,000 patient samples per batch

We also explored what number of barcodes would enable 1,000,000 samples per batch under the second decoder scheme with $k_1 = k_2 = 3$ for scenario 5. A choice of $m_1 = m_2 = 192$ does not suffice, but $m_1 = m_2 = 384$ is sufficient with either $k'_{12} = 5$ or $k'_{12} = 6$. This suggests that we can efficiently test $100,000$ patient samples with a total of $384 + 384 + 10 = 778$ barcodes.

| Scenario | $m_1, m_2$ | $k_1, k_2$ | $k'_{12}$ | Average FNP | Average FPP |
|---|---|---|---|---|---|
| 5 | 192 | 3 | 6 | 0.000232 | 0.0063356 |
| 5 | 384 | 3 | 6 | 0.000344 | 0.0000191 |
| 5 | 384 | 3 | 5 | 0.000016 | 0.0002789 |

Table 2: FPP/FNP with $b = 1,000,000$, $m_3 = 10$, and $p = 0.01$ over 500 iterations.

### 6 Sample Skewing with Template Switching

Sample skewing errors could also occur, as discussed in Supplementary Note 1. This variation could lead to over-representation of some positive samples, preventing detection of samples with lower viral abundance and giving rise to false negatives. We model this in the same way as Supplementary Note 1, adding the possibility of template switching.

#### 6.1 Modeling Continuous Template Switching

The amount of template switching between two barcodes should be dependent on the number of molecules of each barcode present in the PCR pool. To incorporate this, we model the probability of forming a possible template switching product A–B as

$$\Pr[\text{A–B}] = 4\Delta_{switch}\sigma\left(\frac{1}{2}\log\left(\text{Abundance}(A)\right)\right)\sigma\left(\frac{1}{2}\log\left(\text{Abundance}(B)\right)\right)$$

where Abundance($X$) is the number of molecules containing barcode X/average number of molecules per positive barcode 1 or 2, and $\sigma(x)$ is the sigmoid function $\frac{1}{1+e^{-x}}$.

This allows changes in template switching propensity for any given barcode 1 − barcode 2 pair while still bounding the overall probability, with a 4-fold max increase. If a particular template switching product did form, the average of mean(barcode A molecules per barcode 2) and mean(barcode B molecules per barcode 1) molecules of A–B were added.

#### 6.2 Numerical Simulations with Sample Skewing

For numerical simulations, we used the "Amplified" and "Saturated" models from Appendix 1, with the same parameter values. We continued to assume that each NextSeq run generates ≈ 18 million reads per sub-batch. In the simulation, a given barcode pair was called as positive if the number of reads for the barcode pair (calculated by relative abundance multiplied by reads per sub-batch) was greater than or equal to a threshold of $t$ reads. Otherwise, it was called as a negative barcode.

Using the optimized parameter values from section 5, we calculated the average FPPs and FNPs over either 500 or 5000 iterations for scenario 5, shown in Table 3 and Table 4. All results are shown with error rates of $\Delta_{synth} = 0, \Delta_{stoch} = 0.05$, and $\Delta_{switch} = 0.02$.

| Batch Size $b$ | $m_1, m_2$ | $k'_{12}$ | Average FNP | Average FPP | Model |
|---|---|---|---|---|---|
| 100,000 | 96 | 6 | 0.318810 | 0.0000391 | Amplified |
| 100,000 | 192 | 5 | 0.318140 | 0.0000032 | Amplified |
| 100,000 | 384 | 5 | 0.317804 | 0.0000001 | Amplified |
| 100,000 | 96 | 6 | 0.000424 | 0.0002046 | Saturated |
| 100,000 | 192 | 5 | 0.000080 | 0.0000216 | Saturated |
| 100,000 | 384 | 5 | 0.000096 | 0.0000004 | Saturated |

Table 3: FPP/FNP with $t = 100$, $m_3 = 10$, $k_1 = k_2 = 3$, and $p = 0.01$ over 5000 iterations.

| Batch Size $b$ | $m_1, m_2$ | $k'_{12}$ | Average FNP | Average FPP | Model |
|---|---|---|---|---|---|
| 1,000,000 | 384 | 5 | 0.665392 | 0.0000029 | Amplified |
| 1,000,000 | 384 | 6 | 0.665584 | 0.0000000 | Amplified |
| 1,000,000 | 384 | 5 | 0.000000 | 0.0002469 | Saturated |
| 1,000,000 | 384 | 6 | 0.000420 | 0.0000146 | Saturated |

Table 4: FPP/FNP with $t = 100$, $m_3 = 10$, $k_1 = k_2 = 3$, and $p = 0.01$ over 500 iterations.

Overall, the "Amplified" model renders these scenarios untenable, while the "Saturated" model maintains reasonable error rates. High FNPs arise in the "Amplified" model due to crowding out of samples with low viral load by samples with high viral load. We expect LAMP-seq to follow the "Saturated" model, suggesting that these scenarios are robust to sample skewing.

However, it is unclear how accurate the "Saturated" model is. The propagation of sample variation through LAMP-seq needs to be further experimentally characterized, in order to refine this model and to guide parameter choice.

### 7 Code Availability

The scripts used to simulate and plot the models described here are available at the Github repository `https://github.com/dbli2000/SARS-CoV2-Bloom-Filter`.

### References


[1] Burton H. Bloom. 1970. *Space/time trade-offs in hash coding with allowable errors.* Commun. ACM 13, 7 (July 1970), 422–426. DOI: https://doi.org/10.1145/362686.362692


### A Math Appendix

#### A.1 Scenario 4

In this scenario, we can view the barcode 2s and barcode 3s as splitting all of the samples into $m_2 \cdot m_3$ separate non-overlapping Bloom filters each with $\frac{b}{m_2 m_3}$ samples. Plugging these parameters into the equations for $FPP_{\Delta k', m_2}$ and $FNP_{\Delta k', m_2}$ from Supplementary Note 1 give the equations above.

#### A.2 Scenario 5

Each sample is now approximately contained in $k_2$ independent Bloom filters, each with $\frac{k_2 b}{m_2 m_3}$ samples, or $\frac{k_1 k_2 n}{m_2 m_3}$ positive barcode pairs. Solving for the error rate of each barcode pair using results from sections C.2.2 and C.2.3 of Supplementary Note 1 gives the equations above.
